# Comprehensive benchmark and architectural analysis of deep learning models for Nanopore sequencing basecalling

**DOI:** 10.1101/2022.05.17.492272

**Authors:** Marc Pagès-Gallego, Jeroen de Ridder

## Abstract

Nanopore-based DNA sequencing relies on basecalling the electric current signal. Basecalling requires neural networks to achieve competitive accuracies. To improve sequencing accuracy further, new models are continuously proposed. However, benchmarking is currently not standardized, and evaluation metrics and datasets used are defined on a per publication basis, impeding progress in the field. To standardize the process of benchmarking, we unified existing benchmarking datasets and defined a rigorous set of evaluation metrics. We benchmarked the latest seven basecaller models and analyzed their deep learning architectures. Our results show that overall Bonito has the best architecture for basecalling. We find, however, that species bias in training can have a large impact on performance. Our comprehensive evaluation of 90 novel architecture demonstrates that different models excel at reducing different types of errors and using RNNs (LSTM) and a CRF decoder are the main drivers of high performing models.

## 1 Introduction

Sequencing of DNA (or RNA) can be achieved by translocating nucleic acids through a protein nanopore. By passing an electric current through the nanopore, a signal is measured that is representative for the chemical nature of the different nucleotides inside the pore. Therefore, capturing this current yields a signal that can be translated into a DNA sequence. In 2014, Oxford Nanopore Technologies (ONT) released the first commercial sequencing devices based on this principle.

Basecalling is the process of translating the raw current signal to a DNA sequence [1]. It is a fundamental step because almost all downstream applications depend on it [2]. Basecalling is a challenging task due to several reasons. First of all, the current signal level does not correspond to a single base, bus is most dominantly influenced by the several nucleotides that reside inside the pore at any given time. Secondly, DNA molecules do not translocate at a constant speed. Therefore, the number of signal measurements is not a good estimate of sequence length. Instead, the detection of signal changes is required to determine that the next base has entered the pore.

To address the basecalling challenge, a wide array of basecallers has been developed both by ONT and the wider scientific community. Basecallers evolved from statistical tests, to Hidden Markov Models (HMMs) and finally to the use of neural networks [3, 4]. Wick et al. (2019) benchmarked *Chiron* [5] and four other ONT basecallers that were being developed at the time: *Albacore, Guppy, Scrappie* and *Flappie. Chiron* used developments in the speech to text translation field as it applied a Convolutional Neural Network (CNN) to extract the features from the raw signal, a Recurrent Neural Network (RNN) to relate such features in a temporal manner, and a Connectionist Temporal Classification (CTC) decoder [6] to avoid having to segment the signal. Since then, many other basecallers have been developed and published by the community (Figure 1a): *Mincall* [7] used a deep CNN with residual connections [8], *Causalcall* [9] used a CNN with causal dilations [10], *SACall* [11] and *CATCaller* [12] used transformers [13] and lite-transformers [14] respectively, *URNano* [15] used a convolutional U-net with integrated RNNs [16], and *Halcyon* [17] used a sequence-to-sequence (Seq2Seq) model with attention [18, 19]. ONT also updated its main basecaller (*Guppy*) with the architecture from *Bonito*, which substituted the CTC decoder with a conditional random field (CRF) decoder [20].

**Figure 1:**
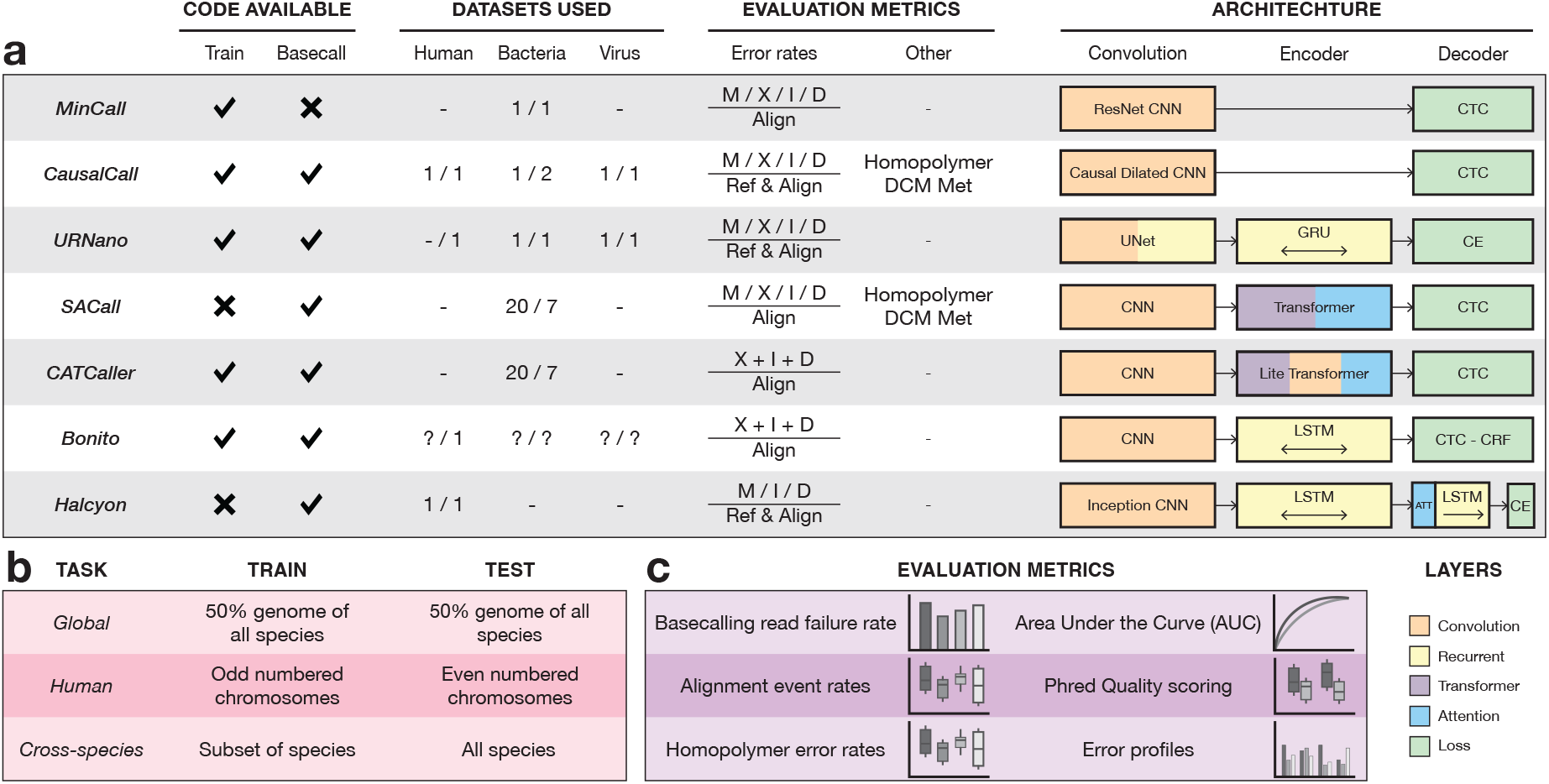
Overview of the latest basecaller models and benchmark. Code available columns indicate whether their github repositories contained scripts to train a model from scratch or basecall using their trained models. Values in the datasets indicate the number of species they used for training (left) and testing (right). Alignment rates reported for: matches (M), mismatches (X), insertions (I) and deletions (D); and whether they were normalized based on the length reference sequence (Ref) or the length of the alignment (Align). Metrics separated by a slash (/) are reported independently, others are reported as a combined error rate. Some models also report homopolymer and methylation site error rates. The architecture section shows a high level overview of the different models. Overview of the different tasks in the benchmark with their respective training and testing data (pink). Overview of the evaluation metrics in the benchmark (purple).

With new AI methods being continuously developed and tailored to various applications, we can reasonably assume that further improvements to basecalling are around the corner. Problematically, there is currently no standardized benchmarking method to evaluate basecalling performance. Instead, evaluation metrics are often defined on a per-publication basis and are limited to only certain aspects of basecalling accuracy (Figure 1a). This may bias conclusions as more favorable metrics are chosen for head to head algorithm comparisons. Furthermore, data used for training and testing is not uniform across publications, which makes it difficult to distinguish between data or neural network architecture driven improvements. This is in stark contrast to e.g., the field of machine learning itself, where new models and methods are benchmarked with community driven evaluation metrics and datasets, such as the CASP competition [21] and the CAMI benchmark [22]. Such efforts facilitate the comparison of new and existing models, and the identification of their respective strengths and weaknesses [23]. It also centralizes all the information, making it easier for users to find the best method for their end goal and lowering the bar to take on the challenge for new developers.

Here we present a wide set of evaluation metrics that can be used to analyze the strengths and weaknesses of basecaller models.

Here we present a comprehensive set of benchmarking tools, including a range of evaluation metrics, that can be used to analyze the strengths and weaknesses of basecaller models. This toolbox can be used as benchmark for the standardized training and cross-comparison of existing and future basecallers. Using this toolbox we benchmarked the latest basecaller models as well as over eighty novel neural network architectures by combining the different components of existing basecaller models. This allowed us to study the influence of the different neural networks components, some of which greatly influence basecalling performance. Finally, we also benchmarked the top performing architectures using different training and testing datasets; allowing us to we evaluate the robustness of the models to training data biases. Together, our work provides insight into what the best model architecture is under what conditions and establishes that model architectures not currently implemented in existing basecalling tools outperform state of the art for certain metrics.

## 2 Results

### 2.1 Benchmark setup

We first developed a basecalling benchmarking framework enabling new and existing basecalling algorithms to be easily compared. Moreover, our benchmark facilitates the study of individual components of basecallers, as different combinations of basecaller components can readily be evaluated. The framework is divided into two main components: (i) standardized datasets for model training and testing (Figure 1b) and (ii) evaluation metrics for extensive assessment of basecalling performance (Figure 1c). The benchmark components are fully automated with minimal dependencies and publicly available on Github, making it easy for developers to focus on algorithm development rather than benchmarking.

#### Datasets and tasks

We gathered datasets that have previously been used for benchmarking, were publicly available for download and used the R9.4 or R9.4.1 pore chemistry. This yielded 615642 reads from 3 human datasets, 1 Lambda phage dataset and 60 bacterial datasets encompassing 26 different bacterial species. After read annotation using *tombo*, 460225 reads (75%) were suitable for benchmarking (Supplemental figure 1; Supplementary Table 1). Because Nanopore sequencing has several possible downstream applications, we defined three different tasks: global, human and cross-species. Each task uses specific portions of training and test data to simulate different scenarios. (i) The **global** task evaluates the performance of a general-purpose model by training and testing the model using data from all available species. In this task, models have access to most data, which is a similar strategy employed in *Causalcall* and *URNano*. (ii) The **human** task evaluates the performance of a human specialized model by training and testing exclusively on human data. This strategy is used in *Halcyon* and *Mincall*, although in the latter *E. coli* is used. (iii) The **cross-species** task evaluates the performance of a trained model on never before seen species. This allows us to evaluate the robustness of the model to overfit on genomic features from the training set, such as k-mer distribution or base modifications. This is achieved by training using a subset of bacterial species and testing on data from all species. This is most similar to the benchmark by Wick et al., (2019) and similar to the evaluation strategy followed in *SACall* and *CATCaller* models. However, here we explicitly take into account the genomic similarity between species when splitting them between train and test sets (see Methods).

#### Evaluation metrics

We define six basecalling evaluation metrics allowing making unbiased and quantitative comparisons of basecalling efficacy across various conditions. (i) Not all reads can be successfully basecalled. Since failed reads are excluded from any subsequent evaluation, it is important to report the **failure rate of the model**. Read basecalling failure falls into one of several categories: not basecalled (the model did not produce any basecalls), not aligned (during evaluation, the alignment algorithm did not produce an alignment), short alignment (the length of the alignment is less than 50% of the length of the reference), or other (other possible errors that prevent the read from being evaluated). (ii) After alignment of a basecalled read to its reference, four types of events can be distinguished: matches, mismatches, insertions and deletions. We report the different types of **alignment rates** independently, since two models may have similar match rates but have different profiles in the types of errors they produce. (iii) **Homopolymers** are particularly difficult to basecall in Nanopore sequencing, for this reason, an independent metric is used to report them. We defined homopolymers as any repetitive sequence of the same base of length 5 or longer. (iv) Basecalling errors can have different distributions depending on the bases that surround the event. **Error profiles**, similar to the trinucleotide context used to define mutational signatures [24], subdivide the errors based on its sequence context to study potential basecalling biases. Ideally, a model’s error profile is unbiased and uniform, for example, to avoid biased variant calling. (v) **PhredQ scores** are values that indicate the confidence of the model in its prediction of a base. Ideally, incorrect calls have low quality scores while high scoring bases are mostly correct. PhredQ scores can be calculated from the predicted distribution of probabilities; however, the calculation of the PhredQ score varies between decoder types, therefore we evaluate the separation of their distributions. (vi) PhredQ scores can also be used to evaluate the model at read level. This is done by evaluating the trade-off between filtering out bad quality reads and the change in performance of the model by calculating the **Area under the Curve (AUC)**.

### 2.2 Benchmark results

To compare existing basecallers and benchmark them in a comparable manner, we re-implemented all their architectures. The results of the human benchmarking challenge are summarized in Figure 2 and Supplementary Table 2.

**Figure 2:**
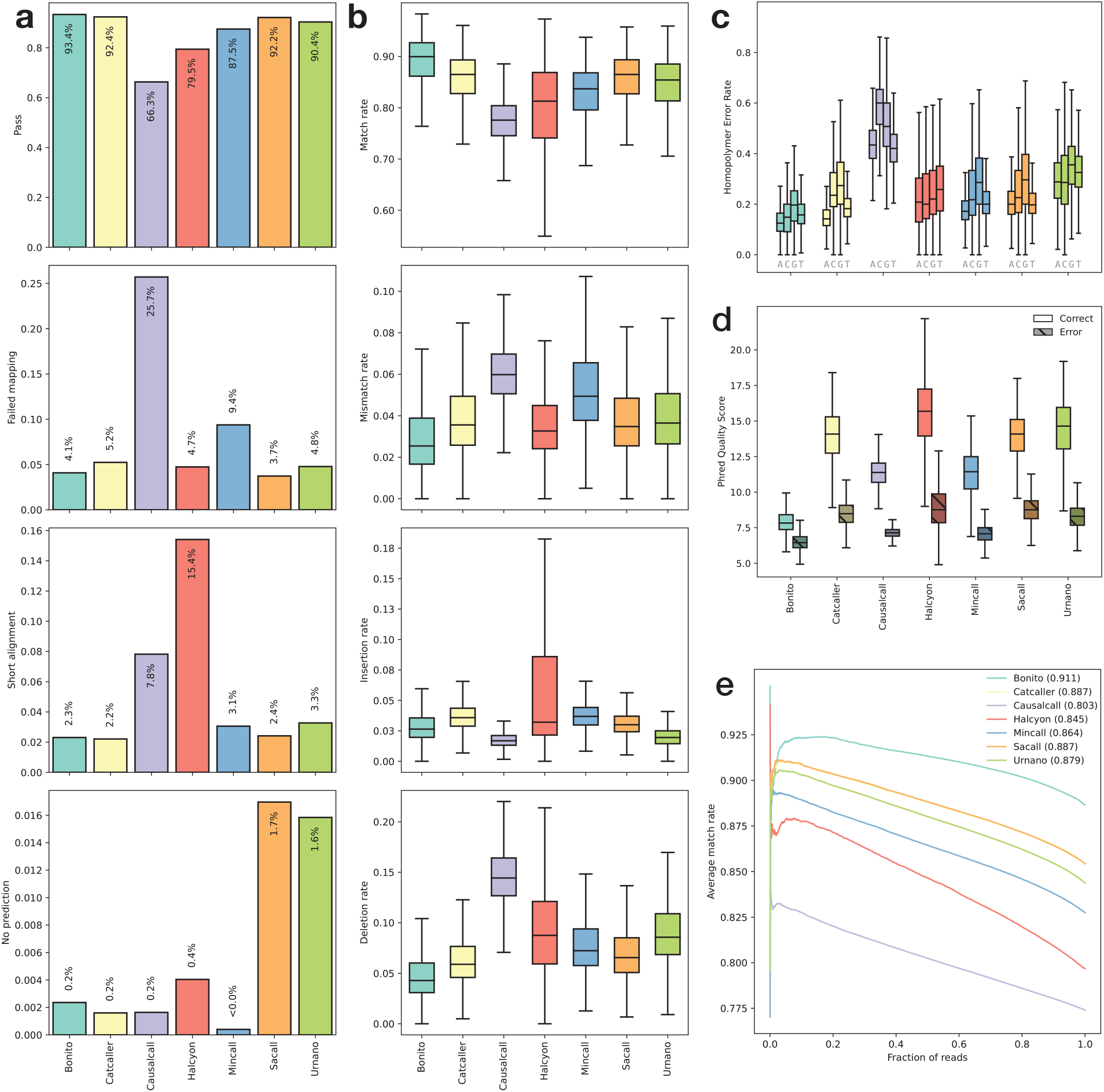
Latest basecallers benchmark on the human task. **(a)** basecalling reads failure rates: pass, failed mapping, short alignment and no prediction (from top top bottom). **(b)** alignment event rates: match, mismatch, insertion and deletion (from top to bottom). **(c)** homopolymer error rates per base. **(d)** PhredQ scores distributions for correctly (light) and incorrectly (dark) predicted bases. **(e)** AUC of match rate sorted by average read PhredQ score.

#### Basecalling failure is a major determinant of reported performance

We first noted that the number of reads that failed basecalling varies substantially. Distinctively, *Causacall, Halcyon* and *Mincall* only managed to properly basecall 66%, 79% and 87% of the reads respectively, while the rest of the models managed >90% (Figure 2a). It is therefore critical to include such a metric in model evaluation since models could be skipping difficult to basecall reads, which would skew results towards a false higher performance.

#### Different methods prevail at different measures

We evaluated the performance of the models using the alignment event rates (Figure 2b). *Bonito* performed best in three out of the four metrics. It has the highest median match rate (90%) and the lowest median mismatch (2.5%) and deletion (4.3%) rates. *Causalcall* achieved the lowest median insertion rate (1.7%); however, it performed worst in the other three metrics with the lowest median match rate (77.6%) and highest mismatch (6%) and deletion (14.4%) rates. *Halcyon* shows the highest variation in performance rates, demonstrating that in addition to the median, the distribution across the reads is important to consider while comparing basecallers. It is therefore critical to not only report the error rates, but also their distributions.

#### Homopolymer error rates are correlated with alignment event rates

Homopolymers are especially difficult to basecall because, for long stretches of the same base, the signal does not change, and since the DNA translocation speed is not constant, the number of measurements is not a good indicator of the length of the homopolymer. For such stretches of DNA, *Bonito* performed best with the lowest median error rate (14.9% averaged across all four bases), and the lowest median error rate for each base individually. *Causalcall* performed significantly worse than the rest of the models with the highest median error rate (44.5% averaged across bases) (Figure 2c). We observed a performance correlation between alignment event rates and homopolymer error rates; however, the latter have a significantly higher error rate likely due to the inherent difficulty of basecalling such stretches.

#### Utility of PhredQ scores varies across methods

To evaluate the relationship between the predicted bases and their PhredQ scores. We first consider the distributions of the scores of the correct and incorrect bases (Figure 2d). For all models, correct bases have higher scores than incorrect bases; *Causalcall* has the smallest overlap between the two distributions (0.7%), followed by the rest of the models with similar overlaps (6-8%) except *Halcyon* (12%) and *Bonito* (32%). Secondly, we calculated AUCs by sorting the reads based on their average quality scores and determining the area under the normalized cumulative score (Figure 2e). All models showed a correlation between read quality and average match rate. Not surprisingly, *Bonito* performed best with an AUC of 0.91. *CATCaller* and *SACAll* are, however, close contenders both with an AUC of 0.886. Importantly, each model has its own PhredQ score offset that determines how their quality scores are calibrated. As a result, quality scores across models, even when compared in a standardized benchmark, are not directly numerically comparable.

#### The signatures of basecalling errors

Finally, we evaluated the different types of mistakes in the context of the two neighboring bases in the basecalls (Supplementary Figure 2a-g). In general, these “error signatures” reveal that there are differences between the accuracy of the models depending on the predicted base context. Across all models, cytosine has low error rates (≈10%) when predicted in CCT or TCT context, however, it can have significantly higher error rates (>30%) when predicted in the context of the 3-mers TCC or TCG. We noticed that many of the contexts with higher error rates contain a CG motif, suggesting the decreased error might be due to the potential methylation status of cytosine. To evaluate if specific models have particular error biases we performed hierarchical clustering on the pairwise Jensen-Shannon divergences between the error signatures (Supplementary Figure 3). This revealed that *Causalcall* and *Halcyon* are the two most different models in terms of “error signatures”. The rest of the models have similar error profiles (lowest Pearson correlation coefficient between them: 0.95). We can conclude that basecalling errors are biased since they are not uniformly distributed across the 3-mer contexts. However, the error profiles are very similar between basecallers, suggesting that training data may play a stronger role in defining these error biases than the architecture of the model itself.

### 2.3 Architecture analysis

The benchmarking setup also allows straightforward investigation of which components of the neural networks provide the main performance gains. To this end, we created novel architectures by combining the convolution, encoder and decoder modules from existing basecallers, as well as some additional modules. We again used the human task to evaluate the different models. In total, ninety different models were evaluated and ranked based on the sum of rankings across all metrics (Figure 3a, Supplementary Figure 5). Out of the original models, *Bonito* again performs best, but reached 9th place in the overall ranking. Consequently, our grid search reveals eight new model architectures that perform better in general. However, improvements in performance made by these models are small; for example in comparison to the *Bonito* model, the alignment event rate improvements and homopolymer error rates are smaller than 1%, suggesting that we may be reaching the performance limits obtainable given the training data used.

**Figure 3:**
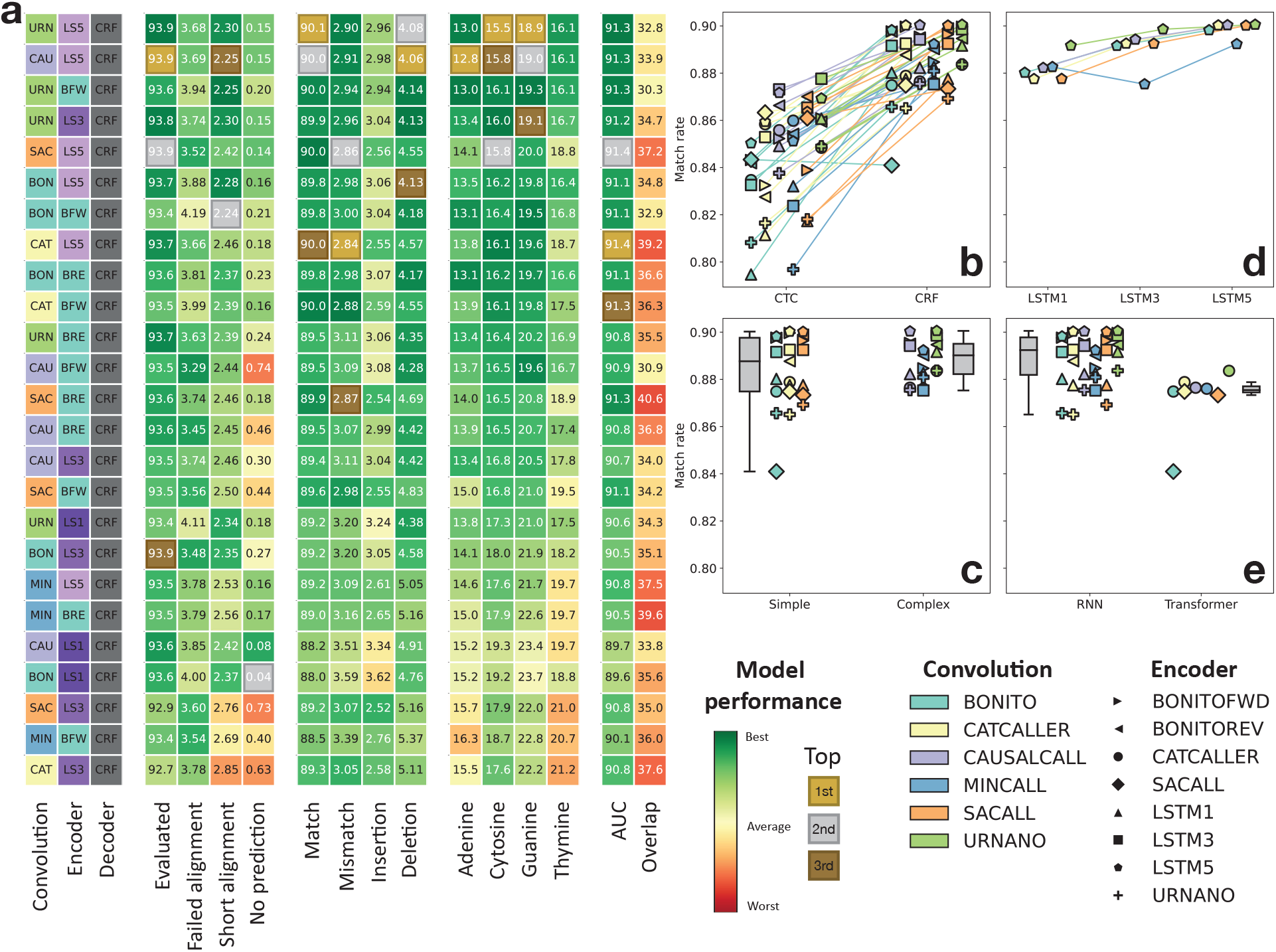
Benchmark of architecture components. **(a)** Top 25 best performing model combinations. **(b)** Comparison of CTC to CRF decoder. **(c)** Comparison of simple (*Bonito, CATCAller, SACall*) to complex (*Causalcall, Mincall, URNano*) convolutions. **(d)** Comparison of bidirection LSTM depth (1, 3 or 5 layers). **(e)** Comparison of RNN to Transformer encoders.

#### CRF decoder is vastly superior to CTC

We observed that most of the high performing models used the CRF decoder module. We therefore compared the change in performance between pairs of models whose only difference was the decoder (Figure 3b, Supplementary Figure 6). We see that for almost all models, using a CRF decoder leads to a general improvement of performance with a mean increase in match rate of 4% and mean decreases of mismatch, insertion and deletion rates of 1%, 1% and 2% respectively. Some exceptions are models that used the *Mincall* or *URNano* convolution modules, which have a mean increase in insertion rates of 1%; although their other alignment rate metrics still improve significantly. This is in concordance with our previous results, where *Causalcall* and *URNano* demonstrated the lowest insertion rates of all the models, showcasing that it is their convolutional architectures that boosts performance for this type of metric. Notably, a decrease in homopolymer error rates is also observed for the models with *Mincall* or *URNano* convolution modules that include a CRF decoder. However, results for the other models are more varied and depend on the base. Consistently with these improvements, we observe an average improvement of 3% in the AUCs. However, the PhredQ overlap between correct and incorrect predictions worsened with a median increase of 30%.

#### Complex convolutions are most robust, but simple convolutions are still very competitive

Another main architectural difference is the complexity and depth of the convolutional layers: ranging from two or three simple convolutional layers like in *Bonito, CATCaller* and *SACall*, to more elaborate convolutional modules like *Causalcall, URNano* or *Mincall*. We find that the top four ranked models use the *URNano* or *Causalcall* architecture (Figure 3a). However, the six following models all use one of the simpler CNNs. More complex convolutional architectures perform better in general, specifically *Causalcall* and *URNano* (Figure 3c, Supplementary Figure 7). Simple convolutional architectures can also perform as good or even better, however they are more dependent on the encoder architecture that follows.

#### RNNs are superior to transformers, and are depth dependent

Transformer layers have gained popularity in other fields due to increased performance and speed [25]. However, our top ten models all use RNN (LSTM) layers in their encoders. A direct comparison shows that RNNs outperform Transformer layers in all the metrics (Figure 3d, Supplementary Figure 8). Therefore the attention mechanism appears to be less useful for this task as the RNNs are able to properly learn the time-axis relationships between samples. We also evaluated whether using deeper LSTM layers has any effect on the performance of the models. We see that there is a clear correlation between depth and performance, encouraging the use of deeper encoders for this task, regardless of the preceding convolutional architecture (Figure 3e, Supplementary Figure 9).

#### Training on multiple species boosts model performance across all models

To assess adaptation of the model to the training dataset, we evaluated the top ten models (ranked based on their performance on the human task) on the global task. We observe that training on multiple species boosts the performance of all ten models across all metrics (Figure 4a). In particular, we observe a 3.2% improvement on mean match rate and 1.1% and 1.3% decrease in mean insertion and deletion rates, respectively (Figure 4b). We also see a general improvement in homopolymer error rates, with decreases of 3.4% (A), 1.5% (C), 2% (G) and 4.2% (T) in mean error rates (Figure 4c). As expected, there is also an improvement in AUCs of 2.9% (Figure 6D). Surprisingly, we observe a significant decrease in the overlap between correct and incorrect base distribution, which was 35% for the human task trained models, and decreased to 21% for the global task trained models (Figure 4c). These improvements in performance could be due to more variance in the data, but it could also be that non-human data is easier to basecall.

**Figure 4:**
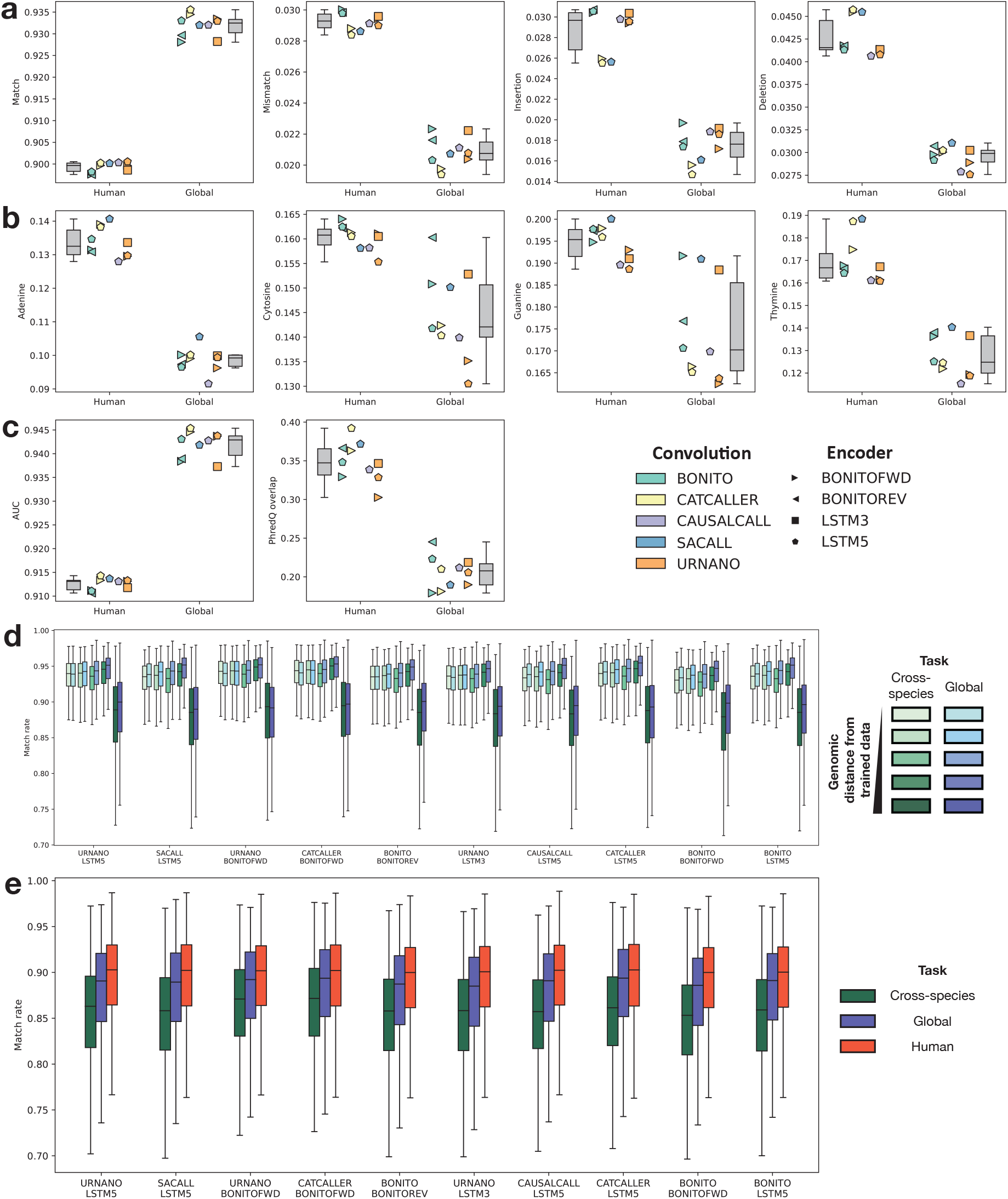
Performance comparison on different tasks. Comparison between the human and global task on the top 10 best performing model combinations: **(a)** Alignment event rates, **(b)** homopolymer error rates and **(c)** PhredQ scoring. **(d)** Comparison between the cross-species (green) and global task (blue). Each boxplot contains a set of species split according to the cross-species task. Darker colors indicate increasing genomic distance from the species in the training set. The darkest colors contain lambda phage and human data. **(e)** Comparison between the human, global and cross-species task only on human data.

#### Lacking a species in the training data penalizes model performance

The cross-species benchmark task is aimed at evaluating the robustness of the models to data from species they have not been trained on. We find that all models performed similarly on all bacterial species, regardless of their k-mer genomics distance to the genomic training set. However, all models perform significantly worse on the two non-bacterial sets (*Homo sapiens* or *Lambda phage*) (Figure 4d). This trend can be seen across all evaluation metrics (Supplementary Figure 10). We therefore asked whether this drop in performance, was due to the lack of training data on those species or whether those particular species are more difficult to basecall. To answer this question, we analyzed the data from the global task in the same manner by splitting the species as in the cross-species task. We observed that, in general, models trained on the global task perform better (0.1%, 0.3%, 0.8%, 0.7% and 0.9% average mean match rate improvement for each set). There is however, a similar trend in which Homo sapiens and Lambda phage reads have a significantly lower performance than the rest of species (Figure 4d). We concluded that, while these species might be more difficult reads in general, the lack of training data on the cross-species task contributed to the lower performance. Finally, we compared the performance of these models only on the *Homo sapiens* data and included the results from the Human task. The results again show that the lack of data is detrimental to the performance of the model (Figure 4e, Supplementary Figure 11). Furthermore, we can observe that all models performed best when only trained with Homo sapiens data, with an average mean match rate improvement of 3.8% and 1.1% when compared to the cross-species and global tasks respectively. This result might encourage the use of species specific models, however, caution should be taken as it is possible that models start to overfit and memorize features like genomic sequences, GC-biases and homopolymer length distributions.

## 3 Discussion

Nanopore basecalling is a critical task in accurate DNA sequencing and currently relies on deep learning, a field in which new algorithms are regularly proposed. However, the lack of consensus in benchmarking data and evaluation metrics makes it complicated and cumbersome to value new contributions by the field. Here we propose a series of tasks and a set of clearly defined evaluation metric swhich can serve as a starting point for what should become a community effort: both by researchers and ONT. In the future both additional tasks and metrics could be added [26]. For example, additional datasets could be added to further study the viability of species-specific models. And computational metrics, such as basecalling speed and compute requirements, could also be added in the future.

Given these proposed tasks and metrics, we benchmarked the latest existing basecallers to evaluate the neural network architectures. To do so, we implemented previously published methods in a coherent framework enabling training on the same data and making a fair comparison not influenced by implementation details or platform. Out of the original models, *Bonito* achieved the best overall performance. However, our results show that other models can obtain better results in other metrics. For this reason, it is important to keep expanding the list of metrics so that end users can choose the best model for their end goals.

To investigate the ideal architecture for basecalling more generally, we created new models by combining their different components. We show that using a CRF decoder, over the more traditional CTC decoder, boosts performance significantly and it is likely the reason why *Bonito* performs so well in the initial benchmark. A CTC decoder predicts each state independently, and it has been its main criticism over the years. For this reason, Seq2Seq models, like Halcyon, were preferred over CTC in other fields since they are able to predict the sequence of states in a conditional manner. However, Seq2seq models are significantly slower than non-recurrent models (CTC and CRF) and require several training steps to be able to deal with large windows of data. The CRF decoder brings the best of both approaches by predicting in a conditional manner while still having a non-recurring decoding step. We also show that deep RNNs (LSTM) are superior to Transformer layers. Transformer layers have gained significant popularity in language processing tasks due to their attention mechanism and speed. However, it appears the attention mechanism is not beneficial for the basecalling task. We hypothesize that, while the attention mechanism might be good for long distance relationships between inputs, the temporal relationships in the electric signal are local enough that RNNs are sufficient for the task. Our results also show that both simple and complex convolutional architectures can achieve competitive performance. Finally, we demonstrate that lack of training data for a particular species decreases model performance; and for species-specific tasks, models trained solely on that particular species have the potential to perform better than more general-purpose models.

## 4 Methodology

### 4.1 Data

We gathered datasets that had previously been used for benchmarking, were available and were sequenced using a R9.4 or R9.4.1 pore chemistry (Supplementary Table 4). With that criteria, we used the bacterial dataset from Wick et al. (2019) and the human genome reference (NA12878/GM12878, Ceph/Utah pedigree) dataset from Jain et al. (2018). The human dataset contained many different sequencing runs. We arbitrarily chose three experiments so that each different ligation kit (rapid, ligation and ultra) would be included: FAB42828, FAF09968 and FAF04090. We also included a Lambda phage dataset that we sequenced. The Lambda phage genome DNA material was purchased from NEB (N3011S). The sample was prepared according to the ONT Ligation KIT and sequenced with a MinION flow cell.

#### Data annotation

Data was annotated using the Tombo resquiggle tool (v1.5.1). First, reads were aligned to the reference sequences using their basecalls. Reference genomes were used as reference sequences for all the datasets; with the exception of the train portion of the bacterial dataset from Wick et al., (2019), which was provided already with a reference sequence for each read. Using Tombo, we aligned the raw signal to the expected signal according to the reference. Reads that did not align to the reference genome or provided a bad resquiggle quality according to Tombo were discarded (Supplementary Figure 1).

#### Tasks definitions

We decided to define three tasks in order to simulate different case scenarios of Nanopore sequencing applications. We simulate this by controlling the data that is used for training and testing in each case. The global task resembles a general purpose model, where the model has been trained on most of the data available. The human task is used to evaluate the feasibility of a species specific model. Finally, the cross-species task is used to evaluate the robustness of the models to unseen species. For each task, we defined a set of reads to be used for training and testing (Supplementary Table 1). In the global task, reads were split according to their mapping position in their respective reference genome (human, bacterial or lambda phage). Bacterial and lambda phage reads that mapped to the first 50% of the genome were used for training, and the rest were used for testing. Reads that mapped to both halves of the genome were discarded. Human reads that mapped odd numbered chromosomes were used for training and reads that mapped to even numbers were used for testing. A total of 100k reads were selected for training and 50k reads for testing. The number of reads was equally split for each species, therefore each species would contribute with a maximum of 3571 for training and 1785 for testing. Some species did not have sufficient reads to reach this quota. A total of 81955 reads were used for training and 47088 reads for testing. For the human task, reads were split as described in the global challenge. For this challenge, reads were also split equally between chromosomes, and a total of 42812 and 25000 reads were used for training and testing. For the cross-species task, species were split between training and testing. First we defined the similarity between species based on the the distribution of 9-mers in their reference genomes. We calculated the pairwise Jensen-Shannon divergence between all species pairs. We then performed single-linkage clustering on the distance matrix and binned the distances into 4 bins (Supplementary Figure 12). We then recursively selected species for training. At each iteration we selected one species and added it to the training pool. We then recalculated the distance between the species in the training pool as a whole, and the rest of the species. We pseudo-randomly added species to the training pool in a manner that, species in the test pool would have a different grades of distance to species in the training pool. We did this by counting the number of test species in each distance bin relative to the training pool, and then picking the species which would change the distribution of counts the least in the next iteration. We continued this process until at least 10 species were chosen for training, with a maximum of 12 allowed. Testing species were divided into different difficulty categories based on their distance to the closest species in the train set (Supplementary Table 5). Similarly to the global task, a total of 50k and 25k reads were selected for training and testing. The testing set was further subdivided between 20k reads, that would come from the test species, and 5k reads, that would come from the train species. Each set of reads was equally divided between the different species. A total of 48013 reads were used for training, 5000 reads were used for testing from the training species, and 24335 reads were used for testing from the testing species.

### 4.2 Models

Model architectures were recreated using Pytorch (v1.9.0) based on their description in their publications and their github repositories. If their implementation was done using Pytorch, code was reused as much as possible (Supplementary Figure 13-Supplementary Figure 27).

#### Model training

Non-recurrent models (all except *Halcyon*) were trained for 5 epochs with a batch size of 64. All models were trained on the same task data which was also given as input in the same order. Reads were sliced in non-overlapping chunks of 2000 data points. Models were trained using an Adam optimizer (initial learning rate = 1e^-^3, *β*_1_ = 0.9, *β*_2_ = 0.999, weight decay = 0). Learning rate was initially increased linearly for 5000 training steps from 0 to the initial learning rate of the optimizer as a warm-up; the learning rate was then decreased using a cosine function until the last training step to a minimum of 1e^-^5. To improve model stability gradients were clipped between −2 and 2. Halcyon was trained similarly to non-recurrent models with the following differences: models were trained first for 1 epoch with non-overlapping chunks of 400 data points, then for 2 epochs with chunks of 1000 data points and finally for 2 epochs with chunks of 2000 data points. This was necessary because training directly using 2000 data points chunks led to unstable model training. This phenomenon is also described in the original Halcyon publication [17], requiting this transfer learning approach to ameliorate the issue. Recurrent models were also trained without warm-up and with a 0.75 scheduled sampling. During training, 5% of the training data was used for validation from which accuracy and loss were calculated without gradients. Validation data was the same for all models. The state of the model was saved every 20000 training steps. The model state was chosen based on the best validation accuracy during training. Models were evaluated on hold out test data from the task being evaluated.

#### Original model recreation and benchmark

*URNano* used cross-entropy as its loss, however, since the objective of the benchmark was basecalling and not signal segmentation, we used a CTC decoder instead. All the other models were recreated as stated in their respective publications, when in doubt their github implementations were used as reference.

#### Comparison of original models to model recreations

Since *Causalcall* and *Halcyon* performed worse than the rest of the models, we evaluated the original models *Causalcall, Halcyon* and *Guppy* model published by the authors, and compared them against our PyTorch implementations (Supplementary Figure 4, Supplementary Table 3). When evaluating the original *Halcyon*, we were unable to completely basecall all 25k reads in the test set due to memory limitations, we therefore compared our recreation based only on the ≈13k reads that were basecalled by the original *Halcyon* model from the human task test set. We used *Guppy* (v5.0.11, latest version) for the comparison between original models and our PyTorch implementations. In terms of reads that we consider evaluable, we see small differences (less than 3%) between the original versions and our implementations of *Causalcall* and *Guppy*. However, we see some differences in the types of failed reads between the original *Causalcall*, which has 7% more reads that failed mapping; whereas our recreation had 3% more reads with short alignments. Surprisingly, basecalls from the original *Halcyon* produce only 46% of reads suitable for evaluation, (30% less than our recreation). A significant 34% of reads failed mapping (30% more than our recreation) to the reference. There is also an increase, although smaller, on the 19% of reads that have short alignments to the reference (5% more than our recreation) (Supplementary Figure 4a). Regarding our implementation of *Guppy*, we find that differences are small, with at most a 2% difference. We then looked at the alignment event rates (Supplementary Figure 4b). Differences between the two *Guppy* models were very small, with the largest being a 1.3% difference in increased match rate from the original version. The original *Causalcall* showed improved match performance, with increased match (2%) and a decreased deletion (6%) rates; however it showed a slight increase in mismatch (1%) and insertion (1%) rates as well as higher variability across reads. Finally, the original *Halcyon* performed worse in all metrics except deletion rate. However, its performances are less variable across reads. Homopolymer error rates show a similar trend (Supplementary Figure 4c), the original *Causalcall* performs significantly better with a more similar to the other models average error rate (25.6%) while the other two models show very similar performances. We finally compared the models regarding their PhredQ scores: when comparing *Bonito* to *Guppy* we saw a large difference in the scale of the scores (Supplementary Figure 4d), however, *Guppy* still had an overlap between the distributions of 28%. On the other hand, the original *Causalcall* showed a significant increase in the overlap between distributions (48%). (Supplementary Figure 4e). Correlating with the event rates results, the original versions of *Causalcall* and *Guppy* performed slightly better than our recreated counterparts with AUCs of 0.837 and 0.937 respectively. The original *Halcyon* does not report any PhredQ scores. With these results, we concluded that although there are some differences between the original models and our recreations, these are minor and could be attributed to training strategies and used data.

#### Architecture analysis

Most models contain a convolutional module that later directly feeds into an encoder (recurrent/transformer) module. To be able to combine modules from different models without changing the original number of channels, we included a linear layer in between the convolution and encoder modules to up-scale or down-scale the number of channels. After this additional linear layer, we applied the last activation function of the preceding convolutional module. Contrary to the other models, the convolution modules from *URNano* and *Causalcall* do not reduce the amount of input timepoints. For those modules, we also included an extra convolution layer with the same configuration as the last convolution layer in *Bonito* (kernel size =19, stride = 5, padding = 9). This layer had the same number of channels as the last convolutional layer of *URNano* or *Causalcall*. This convolution layer was necessary in order to both use transformer encoders and/or a CRF decoder due to memory requirements. We also included three non-used encoder architectures: either one, three or five RNN-LSTM bidirectional stacked layers with 256 channels each.

### 4.3 Evaluation metrics

Evaluation metrics are based on the alignment between the predicted sequence and the reference sequence. Alignment is done using Minimap2 (2.21) [27] with the ONT configuration for all metrics except accuracy. Accuracy is based on the Needleman-Wunsch global alignment algorithm implemented in Parasail (1.2.4) [28]. The global alignment is configured with a match score of 2, a mismatch penalty of 1, a gap opening penalty of 8 and a gap extension penalty of 4. Accuracy is used to evaluate the best performing state of the models during training based on the validation fraction of the data. During training, short sequences have to be aligned, however, during testing, complete reads have to be aligned, for which Minimap2 is necessary.

#### Accuracy

Accuracy is defined as the number of matched bases in the alignment divided by the total number of bases in the reference sequence.

#### Alignment rates

Match, mismatch, insertion and deletion rates are calculated as the number of events of each case divided by the length of the reference unless otherwise stated.

#### Homopolymer error rates

Homopolymer regions are defined as consecutive sequences of the same base of length 5 or longer. Error rates on homopolymer regions are calculated by counting the number of homopolymers with errors (one or more mismatches, insertions or deletions) and dividing it by the number of homopolymer bases.

#### PhredQ scoring

PhredQ scores are calculated using the fast_ctc_decode library from ONT. Average quality scores are calculated for all the correct and incorrect bases for each read. Differences between mean scores between correct and incorrect bases are reported. AUCs are calculated by sorting the basecalled reads according to their mean Phred quality score and calculating the average match rate for cumulative fraction of reads in steps of 50.

#### Error profiles

Error profiles are calculated for all 3-mers by counting the number of events (mismatches for each base, insertions and deletions) in the context of the two neighboring bases of the event itself according to the basecalls. Rates for each event are calculated by dividing each event count by the total number of 3-mer occurrences in the read. Error profiles are also calculated for each base, independently of their context. Randomness of error is defined as the Jensen-Shannon divergence between each 3-mer error profile and their corresponding base error profile.

## 5 Acknowledgements

We thank Tobias Dansen for critical reading of the manuscript. We thank Carlo Vermeulen for critical reading of the manuscript and contribution of the Lambda phage sequencing data. We thank Mike Vella for critical reading of the manuscript. We thank Vlado Menkovski for very helpful discussions.

## 6 Ethics declarations

### Competing interests

Mike Vella works at Oxford Nanopore Technologies, he had no influence on the design or conclusions of the analysis. JdR is cofounder of Cyclomics. JdR has received reimbursement of travel and accommodation expenses to speak at meetings organized by Oxford Nanopore Technologies.

## Notes

### Competing Interest Statement

The authors have declared no competing interest.

### Summary of Updates

Updated abstract and title. Reduced the size of the main text as well as moved one section to the methods. Added additional contributors.

