## Supplemental material for "Comprehensive benchmark and architectural analysis of deep learning models for Nanopore sequencing basecalling"

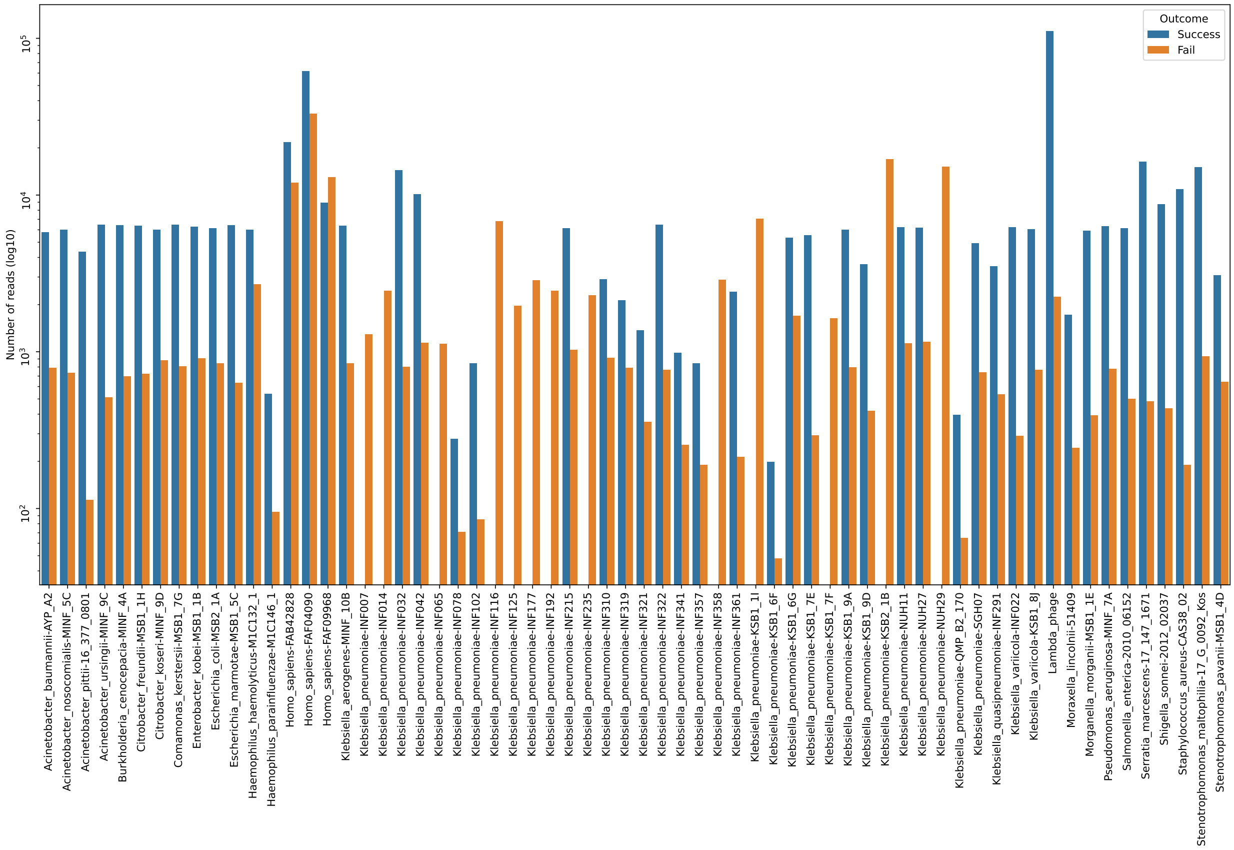

**Supplementary Figure 1: Summary of dataset sizes.** Number of reads per dataset that were successfully resquiggled (blue) or that failed (orange).

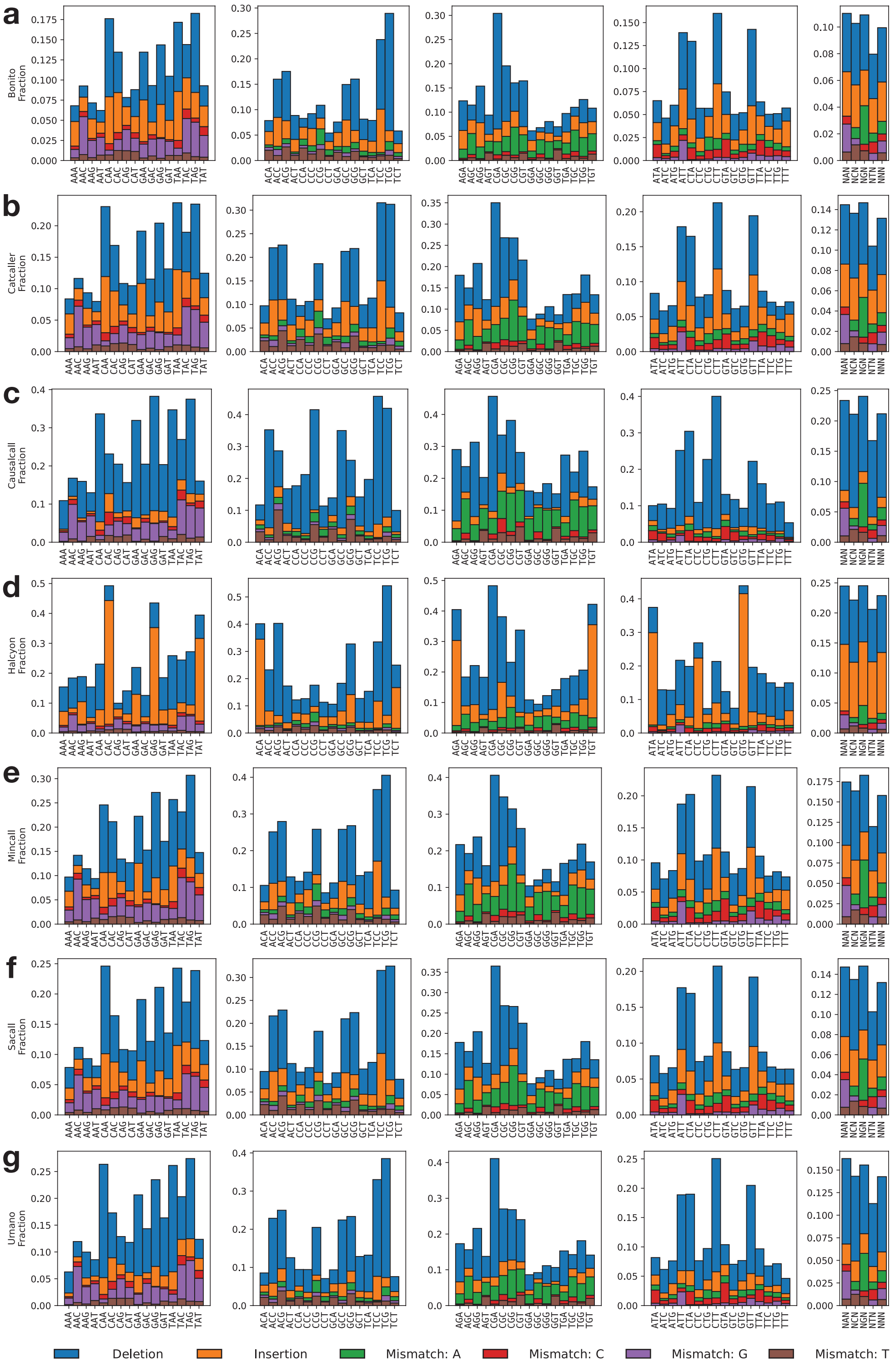

**Supplementary Figure 2: Error profiles of existing basecallers.** Error profiles for each benchmarked model. **(a)** Bonito, **(b)** CATCaller, **(c)** Causalcall, **(d)** Halcyon, **(e)** Mincall, **(f)** SACall and **(g)** URNano. Right most panels show error types aggregated by base regardless of context (NAN, NCN, NGN, NTN) or error types regardless of base (NNN).

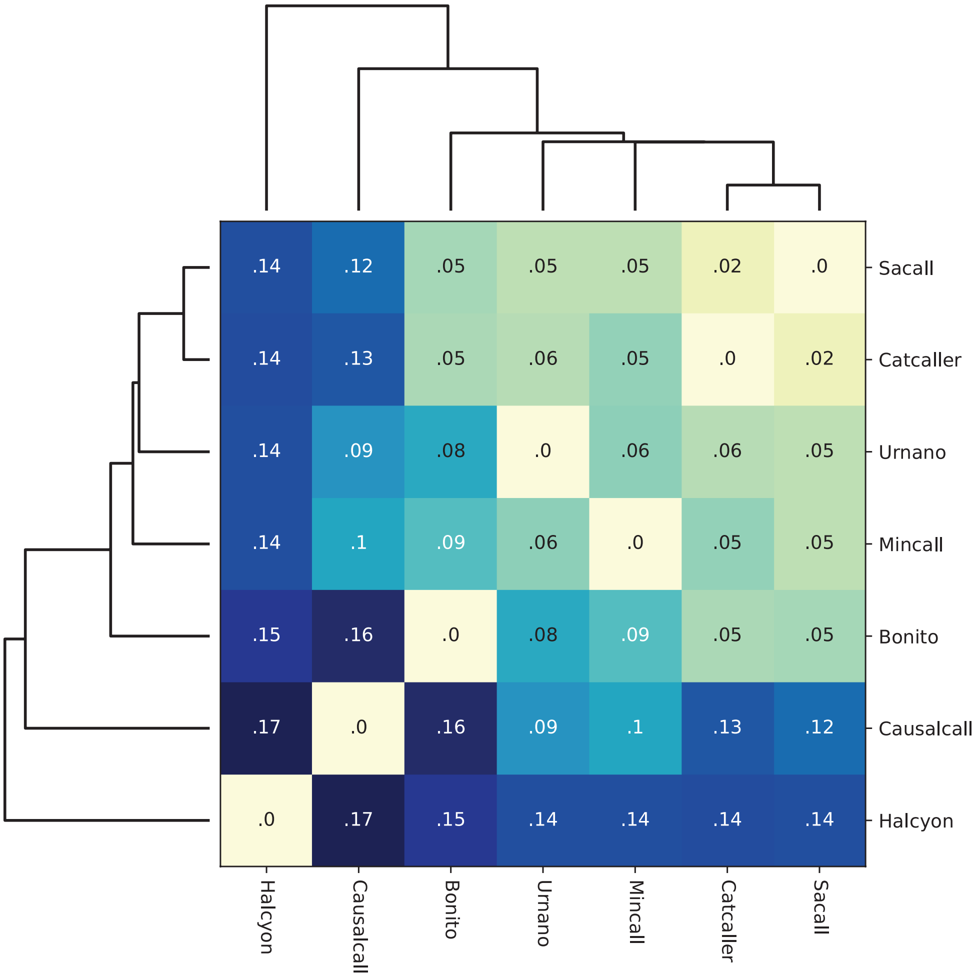

**Supplementary Figure 3: Clustering of error profiles.** Hiearchical clustering of the error profiles from the benchmarked basecallers. Values indicate the Jensen-Shannon divergence between profiles.

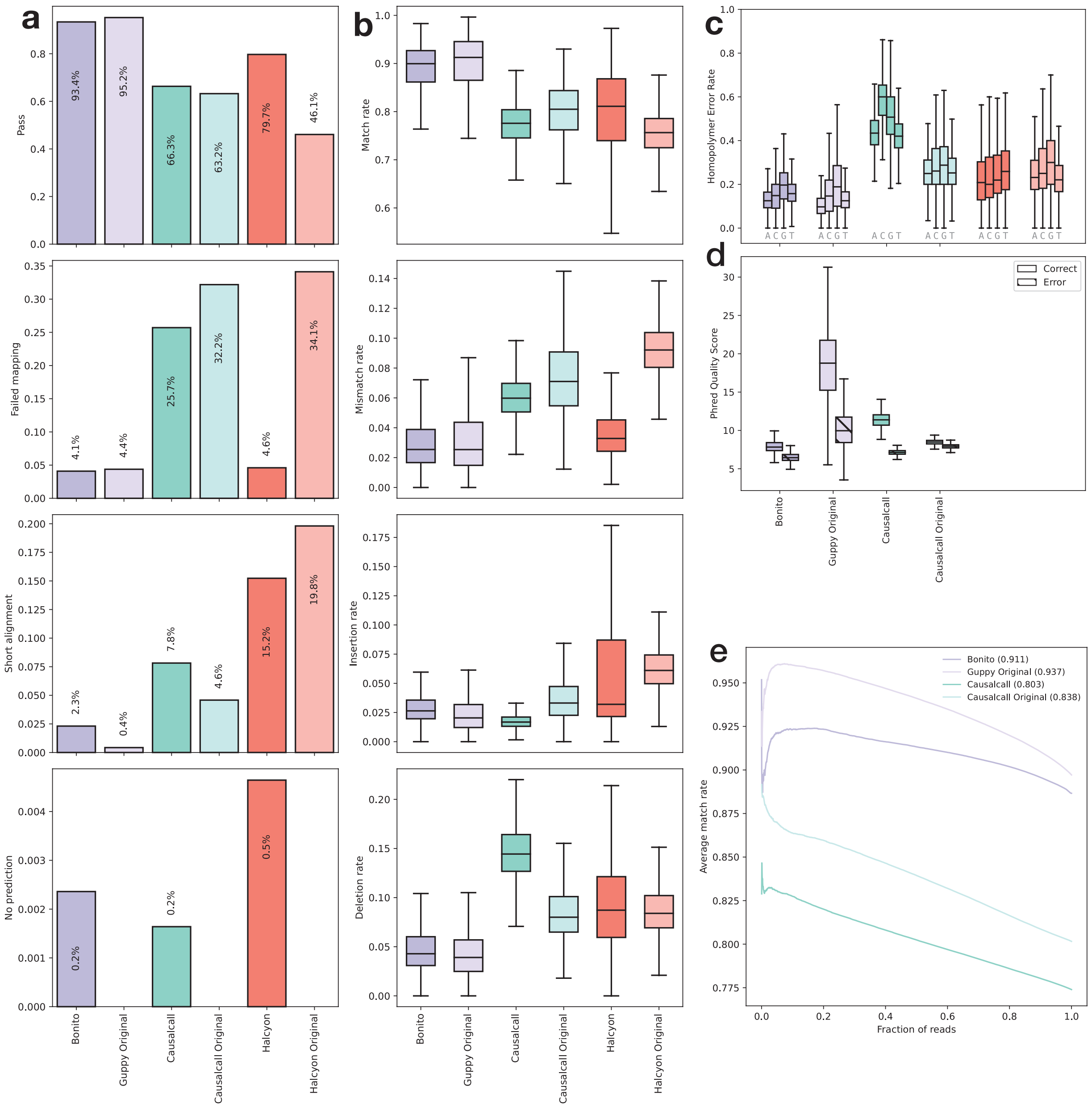

**Supplementary Figure 4: Performance comparison of recreation and original models.** Benchmark of our of *Guppy*, *Causalcall*, *Halcyon* recreations and their original counterparts on the human task. **(a)** basecalling reads failure rates: pass, failed mapping, short alignment and no prediction (from top top bottom). **(b)** alignment event rates: match, mismatch, insertion and deletion (from top to bottom). **(c)** homopolymer error rates per base. **(d)** PhredQ scores distributions for correctly (light) and incorrectly (dark) predicted bases. **(e)** AUC of match rate sorted by average read PhredQ score.

| Convolution | Encoder | Decoder | Evaluated | Failed alignment | Short alignment | No prediction | Match | Mismatch | Insertion | Deletion | Adenine | Cytosine | Guanine | Thymine | AUC | PhredQ overlap |
|---|---|---|---|---|---|---|---|---|---|---|---|---|---|---|---|---|
| URN | LS5 | CRF | 93.9 | 3.68 | 2.30 | 0.15 | 90.1 | 2.90 | 2.96 | 4.08 | 13.0 | 15.5 | 18.9 | 16.1 | 91.3 | 32.8 |
| CAU | LS5 | CRF | 93.9 | 3.69 | 2.25 | 0.15 | 90.0 | 2.91 | 2.98 | 4.06 | 12.8 | 15.8 | 19.0 | 16.1 | 91.3 | 33.9 |
| URN | BFW | CRF | 93.6 | 3.94 | 2.25 | 0.20 | 90.0 | 2.94 | 2.94 | 4.14 | 13.0 | 16.1 | 19.3 | 16.1 | 91.3 | 30.3 |
| URN | LS3 | CRF | 93.8 | 3.74 | 2.30 | 0.15 | 89.9 | 2.96 | 3.04 | 4.13 | 13.4 | 16.0 | 19.1 | 16.7 | 91.2 | 34.7 |
| SAC | LS5 | CRF | 93.9 | 3.52 | 2.42 | 0.14 | 90.0 | 2.86 | 2.56 | 4.55 | 14.1 | 15.8 | 20.0 | 18.8 | 91.4 | 37.2 |
| BON | LS5 | CRF | 93.7 | 3.88 | 2.28 | 0.16 | 89.8 | 2.98 | 3.06 | 4.13 | 13.5 | 16.2 | 19.8 | 16.4 | 91.1 | 34.8 |
| BON | BFW | CRF | 93.4 | 4.19 | 2.24 | 0.21 | 89.8 | 3.00 | 3.04 | 4.18 | 13.1 | 16.4 | 19.5 | 16.8 | 91.1 | 32.9 |
| CAT | LS5 | CRF | 93.7 | 3.66 | 2.46 | 0.18 | 90.0 | 2.84 | 2.55 | 4.57 | 13.8 | 16.1 | 19.6 | 18.7 | 91.4 | 39.2 |
| BON | BRE | CRF | 93.6 | 3.81 | 2.37 | 0.23 | 89.8 | 2.98 | 3.07 | 4.17 | 13.1 | 16.2 | 19.7 | 16.6 | 91.1 | 36.6 |
| CAT | BFW | CRF | 93.5 | 3.99 | 2.39 | 0.16 | 90.0 | 2.88 | 2.59 | 4.55 | 13.9 | 16.1 | 19.8 | 17.5 | 91.3 | 36.3 |
| URN | BRE | CRF | 93.7 | 3.63 | 2.39 | 0.24 | 89.5 | 3.11 | 3.06 | 4.35 | 13.4 | 16.4 | 20.2 | 16.9 | 90.8 | 35.5 |
| CAU | BFW | CRF | 93.5 | 3.29 | 2.44 | 0.74 | 89.5 | 3.09 | 3.08 | 4.28 | 13.7 | 16.5 | 19.6 | 16.7 | 90.9 | 30.9 |
| SAC | BRE | CRF | 93.6 | 3.74 | 2.46 | 0.18 | 89.9 | 2.87 | 2.54 | 4.69 | 14.0 | 16.5 | 20.8 | 18.9 | 91.3 | 40.6 |
| CAU | BRE | CRF | 93.6 | 3.45 | 2.45 | 0.46 | 89.5 | 3.07 | 2.99 | 4.42 | 13.9 | 16.5 | 20.7 | 17.4 | 90.8 | 36.8 |
| CAU | LS3 | CRF | 93.5 | 3.74 | 2.46 | 0.30 | 89.4 | 3.11 | 3.04 | 4.42 | 13.4 | 16.8 | 20.5 | 17.8 | 90.7 | 34.0 |
| SAC | BFW | CRF | 93.5 | 3.56 | 2.50 | 0.44 | 89.6 | 2.98 | 2.55 | 4.83 | 15.0 | 16.8 | 21.0 | 19.5 | 91.1 | 34.2 |
| URN | LS1 | CRF | 93.4 | 4.11 | 2.34 | 0.18 | 89.2 | 3.20 | 3.24 | 4.38 | 13.8 | 17.3 | 21.0 | 17.5 | 90.6 | 34.3 |
| BON | LS3 | CRF | 93.9 | 3.48 | 2.35 | 0.27 | 89.2 | 3.20 | 3.05 | 4.58 | 14.1 | 18.0 | 21.9 | 18.2 | 90.5 | 35.1 |
| MIN | LS5 | CRF | 93.5 | 3.78 | 2.53 | 0.16 | 89.2 | 3.09 | 2.61 | 5.05 | 14.6 | 17.6 | 21.7 | 19.7 | 90.8 | 37.5 |
| MIN | BRE | CRF | 93.5 | 3.79 | 2.56 | 0.17 | 89.0 | 3.16 | 2.65 | 5.16 | 15.0 | 17.9 | 22.6 | 19.7 | 90.5 | 39.6 |
| CAU | LS1 | CRF | 93.6 | 3.85 | 2.42 | 0.08 | 88.2 | 3.51 | 3.34 | 4.91 | 15.2 | 19.3 | 23.4 | 19.7 | 89.7 | 33.8 |
| BON | LS1 | CRF | 93.6 | 4.00 | 2.37 | 0.04 | 88.0 | 3.59 | 3.62 | 4.76 | 15.2 | 19.2 | 23.7 | 18.8 | 89.6 | 35.6 |
| SAC | LS3 | CRF | 92.9 | 3.60 | 2.76 | 0.73 | 89.2 | 3.07 | 2.52 | 5.16 | 15.7 | 17.9 | 22.0 | 21.0 | 90.8 | 35.0 |
| MIN | BFW | CRF | 93.4 | 3.54 | 2.69 | 0.40 | 88.5 | 3.39 | 2.76 | 5.37 | 16.3 | 18.7 | 22.8 | 20.7 | 90.1 | 36.0 |
| CAT | LS3 | CRF | 92.7 | 3.78 | 2.85 | 0.63 | 89.3 | 3.05 | 2.58 | 5.11 | 15.5 | 17.6 | 22.2 | 21.2 | 90.8 | 37.6 |
| URN | URN | CRF | 93.1 | 3.86 | 2.72 | 0.27 | 88.4 | 3.46 | 3.14 | 5.02 | 15.5 | 19.0 | 23.3 | 18.9 | 89.9 | 36.9 |
| CAT | BRE | CRF | 92.7 | 4.42 | 2.74 | 0.12 | 88.8 | 3.07 | 2.46 | 5.69 | 16.9 | 18.3 | 24.1 | 20.9 | 90.4 | 43.3 |
| CAU | CAT | CTC | 91.3 | 6.08 | 2.44 | 0.18 | 85.6 | 4.29 | 4.99 | 5.12 | 13.3 | 17.6 | 19.8 | 15.6 | 87.8 | 11.9 |
| URN | CAT | CRF | 92.7 | 4.18 | 2.69 | 0.44 | 88.4 | 3.46 | 3.16 | 5.01 | 15.8 | 19.8 | 23.3 | 19.4 | 89.8 | 35.2 |
| MIN | URN | CRF | 93.2 | 3.88 | 2.82 | 0.10 | 88.1 | 3.43 | 2.95 | 5.52 | 16.5 | 20.6 | 26.2 | 20.9 | 89.9 | 43.0 |
| SAC | LS3 | CTC | 92.0 | 4.66 | 2.45 | 0.94 | 86.5 | 3.92 | 3.73 | 5.79 | 14.6 | 19.2 | 21.2 | 18.1 | 89.0 | 8.20 |
| BON | SAC | CTC | 91.0 | 6.52 | 2.32 | 0.13 | 84.3 | 4.69 | 5.57 | 5.38 | 13.6 | 18.4 | 21.8 | 15.3 | 86.5 | 9.14 |
| CAT | LS1 | CRF | 93.4 | 3.91 | 2.67 | 0.07 | 87.8 | 3.54 | 2.90 | 5.79 | 18.0 | 21.4 | 28.5 | 23.9 | 89.5 | 34.3 |
| MIN | CAT | CTC | 91.8 | 5.85 | 2.27 | 0.08 | 86.0 | 4.17 | 3.75 | 6.07 | 14.7 | 21.2 | 24.4 | 17.5 | 88.4 | 8.13 |
| SAC | BRE | CTC | 92.2 | 4.19 | 2.43 | 1.18 | 87.0 | 3.76 | 3.77 | 5.45 | 14.8 | 19.3 | 21.8 | 19.8 | 89.5 | 11.2 |
| MIN | LS1 | CRF | 92.9 | 3.94 | 2.84 | 0.32 | 88.3 | 3.40 | 2.84 | 5.49 | 16.5 | 20.2 | 26.0 | 22.3 | 89.9 | 37.1 |
| BON | LS5 | CTC | 90.9 | 6.00 | 2.65 | 0.41 | 85.0 | 4.50 | 5.40 | 5.06 | 13.0 | 17.1 | 21.2 | 15.8 | 87.4 | 9.06 |
| SAC | LS1 | CRF | 93.2 | 4.08 | 2.63 | 0.05 | 87.8 | 3.56 | 2.99 | 5.68 | 17.7 | 21.9 | 27.9 | 23.3 | 89.5 | 35.9 |
| SAC | LS5 | CTC | 91.5 | 5.38 | 2.63 | 0.48 | 86.4 | 3.99 | 3.97 | 5.67 | 14.2 | 20.0 | 21.4 | 18.3 | 88.9 | 8.35 |
| BON | BFW | CTC | 90.5 | 6.64 | 2.62 | 0.24 | 84.3 | 4.85 | 5.58 | 5.24 | 13.4 | 18.0 | 21.3 | 16.4 | 86.7 | 7.96 |
| BON | CAT | CTC | 90.0 | 7.48 | 2.32 | 0.18 | 83.5 | 5.01 | 5.67 | 5.84 | 13.7 | 17.8 | 21.5 | 14.7 | 85.5 | 7.96 |
| CAU | URN | CRF | 92.6 | 4.64 | 2.52 | 0.24 | 87.5 | 3.73 | 3.53 | 5.18 | 15.5 | 20.2 | 26.0 | 19.9 | 89.3 | 41.4 |
| BON | CAT | CRF | 93.3 | 4.04 | 2.59 | 0.05 | 87.5 | 3.79 | 3.68 | 5.04 | 16.2 | 21.2 | 28.1 | 21.5 | 88.4 | 46.2 |
| CAT | LS3 | CTC | 91.4 | 5.58 | 2.38 | 0.60 | 85.3 | 4.40 | 4.39 | 5.90 | 14.5 | 21.2 | 21.9 | 17.7 | 87.8 | 7.12 |
| CAU | SAC | CTC | 90.7 | 6.73 | 2.37 | 0.22 | 84.2 | 4.96 | 5.09 | 5.78 | 14.0 | 20.2 | 22.7 | 15.8 | 86.3 | 8.10 |
| CAU | CAT | CRF | 92.8 | 4.56 | 2.48 | 0.18 | 87.6 | 3.73 | 3.33 | 5.28 | 17.4 | 22.0 | 27.1 | 20.3 | 88.6 | 40.9 |

| Convolution | Encoder | Decoder | Evaluated | Failed alignment | Short alignment | No prediction | Match | Mismatch | Insertion | Deletion | Adenine | Cytosine | Guanine | Thymine | AUC | PhredQ overlap |
|---|---|---|---|---|---|---|---|---|---|---|---|---|---|---|---|---|
| BON | LS3 | CTC | 89.4 | 7.42 | 2.68 | 0.52 | 83.3 | 5.25 | 6.10 | 5.37 | 13.2 | 19.0 | 20.6 | 15.4 | 85.6 | 7.47 |
| SAC | CAT | CTC | 92.1 | 5.47 | 2.32 | 0.15 | 86.0 | 4.07 | 3.77 | 6.16 | 16.0 | 25.2 | 29.4 | 20.2 | 88.4 | 8.00 |
| CAT | LS5 | CTC | 91.2 | 5.67 | 2.69 | 0.45 | 85.9 | 4.16 | 3.87 | 6.02 | 14.6 | 20.7 | 23.3 | 18.5 | 88.5 | 8.05 |
| CAT | CAT | CRF | 92.7 | 4.39 | 2.77 | 0.14 | 87.9 | 3.49 | 3.04 | 5.58 | 17.1 | 23.6 | 30.1 | 27.0 | 89.1 | 49.0 |
| MIN | BFW | CTC | 91.6 | 4.72 | 2.50 | 1.20 | 85.3 | 4.50 | 3.75 | 6.40 | 15.0 | 20.2 | 23.6 | 19.4 | 87.8 | 5.57 |
| MIN | CAT | CRF | 92.7 | 4.30 | 2.87 | 0.08 | 87.6 | 3.55 | 2.95 | 5.89 | 18.7 | 23.3 | 30.0 | 23.3 | 89.0 | 44.9 |
| URN | LS3 | CTC | 91.9 | 4.81 | 2.92 | 0.33 | 87.8 | 3.04 | 1.75 | 7.43 | 26.3 | 24.8 | 30.3 | 30.9 | 90.3 | 10.4 |
| MIN | LS3 | CRF | 92.4 | 3.76 | 2.90 | 0.90 | 87.5 | 3.68 | 2.83 | 5.95 | 18.1 | 21.2 | 26.8 | 22.8 | 89.3 | 36.6 |
| MIN | LS5 | CTC | 91.1 | 5.63 | 2.67 | 0.60 | 85.1 | 4.54 | 3.91 | 6.42 | 14.6 | 21.1 | 22.9 | 17.3 | 87.6 | 7.99 |
| URN | CAT | CTC | 90.5 | 6.36 | 2.84 | 0.26 | 84.8 | 4.67 | 4.78 | 5.70 | 14.2 | 19.1 | 21.6 | 16.9 | 87.2 | 10.8 |
| BON | BRE | CTC | 90.1 | 6.54 | 2.86 | 0.54 | 84.2 | 4.88 | 5.54 | 5.39 | 13.5 | 18.1 | 21.5 | 16.6 | 86.6 | 12.8 |
| MIN | BRE | CTC | 91.6 | 4.74 | 2.59 | 1.03 | 85.4 | 4.47 | 3.91 | 6.18 | 14.5 | 20.7 | 23.3 | 19.4 | 88.0 | 10.3 |
| CAU | LS5 | CTC | 91.7 | 2.59 | 2.96 | 2.71 | 87.2 | 3.17 | 1.72 | 7.93 | 26.9 | 27.4 | 30.2 | 31.6 | 89.8 | 9.64 |
| CAT | SAC | CTC | 92.1 | 4.95 | 2.38 | 0.55 | 86.3 | 3.93 | 3.35 | 6.38 | 18.7 | 25.6 | 28.9 | 22.8 | 88.8 | 8.15 |
| CAU | LS3 | CTC | 91.7 | 3.65 | 2.91 | 1.71 | 87.3 | 3.13 | 1.68 | 7.87 | 25.6 | 29.4 | 32.2 | 31.5 | 89.9 | 9.62 |
| SAC | SAC | CTC | 91.9 | 3.44 | 2.45 | 2.20 | 86.1 | 3.82 | 3.01 | 7.07 | 22.4 | 26.6 | 31.3 | 25.0 | 88.6 | 9.23 |
| URN | LS5 | CTC | 91.7 | 2.91 | 3.01 | 2.40 | 86.9 | 3.35 | 1.88 | 7.83 | 28.0 | 26.8 | 32.3 | 31.6 | 89.5 | 9.96 |
| MIN | SAC | CTC | 90.9 | 6.61 | 2.16 | 0.36 | 83.4 | 5.20 | 4.78 | 6.63 | 16.1 | 22.2 | 27.9 | 17.6 | 85.7 | 5.67 |
| SAC | SAC | CRF | 92.8 | 3.61 | 2.54 | 1.08 | 87.3 | 3.73 | 3.43 | 5.50 | 20.1 | 24.7 | 30.7 | 29.9 | 88.6 | 47.3 |
| CAT | CAT | CTC | 92.0 | 5.32 | 2.38 | 0.32 | 85.8 | 4.09 | 3.65 | 6.41 | 16.3 | 28.0 | 30.6 | 20.2 | 88.3 | 10.1 |
| BON | URN | CTC | 84.4 | 12.3 | 3.18 | 0.14 | 80.8 | 6.29 | 7.09 | 5.79 | 13.2 | 19.0 | 22.2 | 15.4 | 83.1 | 8.85 |
| SAC | URN | CRF | 92.3 | 4.52 | 2.89 | 0.26 | 86.9 | 3.83 | 3.28 | 5.96 | 17.6 | 22.5 | 29.0 | 24.2 | 88.9 | 45.8 |
| CAU | BRE | CTC | 91.4 | 2.91 | 3.03 | 2.70 | 86.6 | 3.49 | 1.98 | 7.89 | 29.3 | 28.1 | 30.8 | 31.6 | 89.3 | 11.5 |
| CAT | SAC | CRF | 92.4 | 4.11 | 2.75 | 0.70 | 87.5 | 3.63 | 3.28 | 5.59 | 20.9 | 24.8 | 30.4 | 29.7 | 88.7 | 48.0 |
| BON | LS1 | CTC | 81.8 | 15.0 | 3.14 | 0.06 | 79.5 | 6.78 | 6.66 | 7.08 | 12.9 | 20.6 | 22.9 | 14.4 | 81.6 | 8.20 |
| BON | URN | CRF | 92.2 | 4.25 | 2.71 | 0.86 | 86.6 | 4.09 | 3.76 | 5.57 | 16.7 | 23.4 | 28.4 | 21.4 | 88.3 | 41.4 |
| MIN | LS1 | CTC | 87.7 | 9.14 | 3.04 | 0.08 | 83.2 | 5.23 | 4.49 | 7.06 | 15.2 | 23.1 | 24.2 | 17.7 | 85.9 | 8.08 |
| URN | BFW | CTC | 91.3 | 2.38 | 3.10 | 3.25 | 86.1 | 3.66 | 2.03 | 8.22 | 26.7 | 29.4 | 32.0 | 33.0 | 88.7 | 9.10 |
| SAC | BFW | CTC | 89.5 | 7.12 | 2.82 | 0.60 | 83.9 | 5.01 | 4.48 | 6.61 | 15.6 | 22.4 | 25.2 | 18.9 | 86.6 | 5.83 |
| CAT | LS1 | CTC | 84.8 | 12.2 | 3.00 | 0.04 | 81.1 | 5.97 | 5.19 | 7.68 | 15.4 | 23.6 | 24.6 | 16.6 | 83.6 | 5.62 |
| URN | BRE | CTC | 91.2 | 3.26 | 3.02 | 2.54 | 85.9 | 3.79 | 2.17 | 8.07 | 26.6 | 28.6 | 32.2 | 34.3 | 88.7 | 8.32 |
| SAC | LS1 | CTC | 85.7 | 11.3 | 2.96 | 0.03 | 81.7 | 5.70 | 5.13 | 7.43 | 14.9 | 22.9 | 25.5 | 16.1 | 84.2 | 9.10 |
| MIN | LS3 | CTC | 88.0 | 9.08 | 2.79 | 0.18 | 82.4 | 5.60 | 4.72 | 7.29 | 15.2 | 23.8 | 26.4 | 17.4 | 84.8 | 7.07 |
| URN | LS1 | CTC | 90.8 | 4.88 | 3.17 | 1.16 | 86.1 | 3.63 | 1.98 | 8.31 | 26.4 | 29.8 | 32.0 | 33.5 | 88.8 | 7.24 |
| CAT | BFW | CTC | 88.8 | 7.95 | 2.88 | 0.39 | 83.2 | 5.31 | 4.76 | 6.67 | 15.9 | 23.3 | 25.1 | 19.6 | 85.9 | 5.71 |
| CAT | URN | CRF | 91.5 | 4.64 | 3.28 | 0.61 | 86.5 | 3.84 | 3.25 | 6.39 | 16.6 | 24.4 | 30.9 | 22.5 | 88.7 | 46.8 |
| BON | SAC | CRF | 91.3 | 3.08 | 2.80 | 2.82 | 84.1 | 5.07 | 5.22 | 5.60 | 19.5 | 21.1 | 24.9 | 27.4 | 86.3 | 38.9 |
| CAT | URN | CTC | 86.9 | 10.2 | 2.82 | 0.10 | 81.6 | 5.88 | 5.48 | 6.99 | 16.1 | 26.1 | 29.0 | 17.7 | 84.1 | 6.08 |
| CAU | BFW | CTC | 89.9 | 3.04 | 3.67 | 3.41 | 85.2 | 3.63 | 1.81 | 9.32 | 32.3 | 30.0 | 34.4 | 36.6 | 88.1 | 8.31 |
| SAC | URN | CTC | 87.3 | 9.78 | 2.76 | 0.11 | 81.8 | 5.82 | 5.06 | 7.31 | 16.7 | 27.8 | 26.3 | 19.0 | 84.3 | 6.79 |
| MIN | URN | CTC | 81.5 | 15.2 | 3.23 | 0.05 | 79.7 | 6.75 | 5.32 | 8.24 | 16.0 | 27.2 | 31.0 | 17.2 | 82.0 | 4.69 |
| CAU | LS1 | CTC | 89.0 | 6.56 | 3.80 | 0.64 | 84.9 | 3.70 | 1.74 | 9.63 | 30.2 | 35.4 | 37.8 | 32.9 | 87.8 | 7.19 |
| URN | URN | CTC | 89.8 | 5.40 | 3.40 | 1.42 | 84.9 | 4.05 | 2.17 | 8.87 | 29.5 | 30.7 | 32.7 | 33.4 | 87.7 | 7.06 |
| CAT | BRE | CTC | 87.9 | 8.14 | 3.03 | 0.94 | 82.8 | 5.50 | 4.82 | 6.91 | 16.4 | 23.7 | 28.1 | 19.5 | 85.5 | 10.1 |
| CAU | URN | CTC | 88.2 | 5.74 | 3.83 | 2.20 | 83.8 | 4.08 | 1.88 | 10.3 | 35.0 | 34.7 | 42.3 | 35.8 | 86.6 | 6.68 |

**Supplementary Figure 5: Benchmark of architecture components.** Performance of all combined components benchmarked on the human task. Models sorted based on their overall performance ranking. Gradient coloring indicates: good performance (green), average performance (yellow), bad performance (red). Top three models per metric are indicated in gold (1st), silver (2nd) and bronze (3rd) colors.

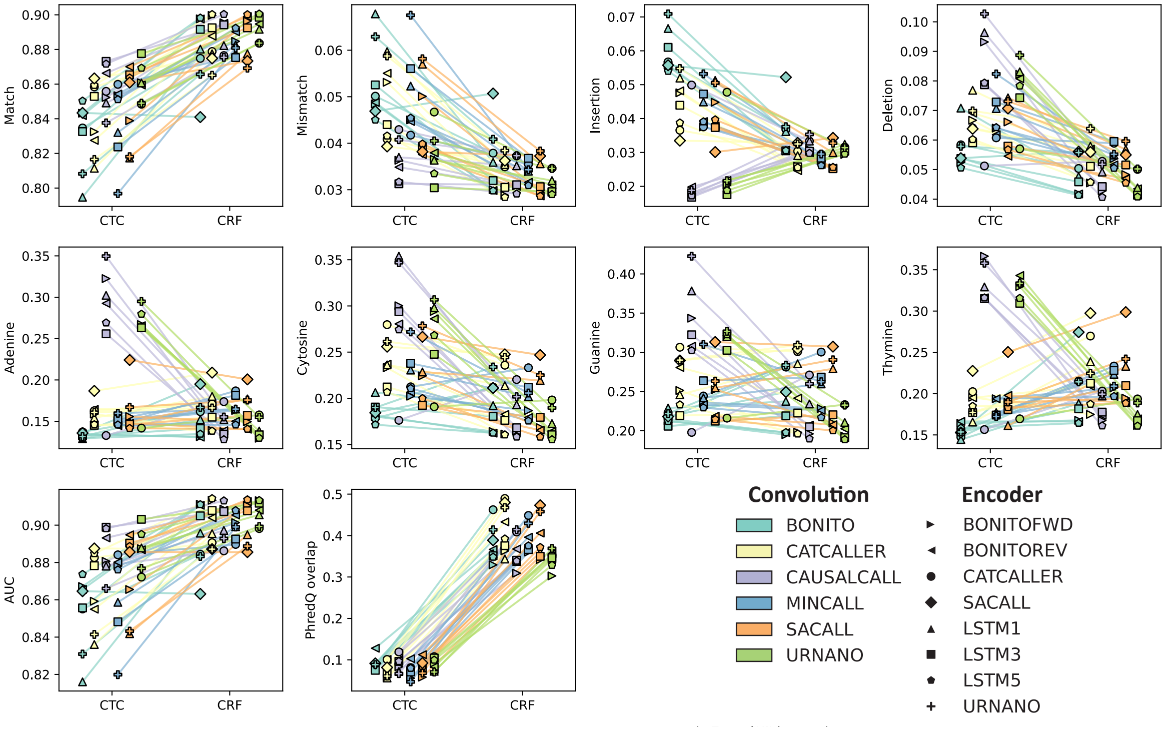

**Supplementary Figure 6: Comparison of CTC and CRF decoders.** Pair-wise performance comparison of models that share the same architecture with the exception of the decoder (CTC or CRF).

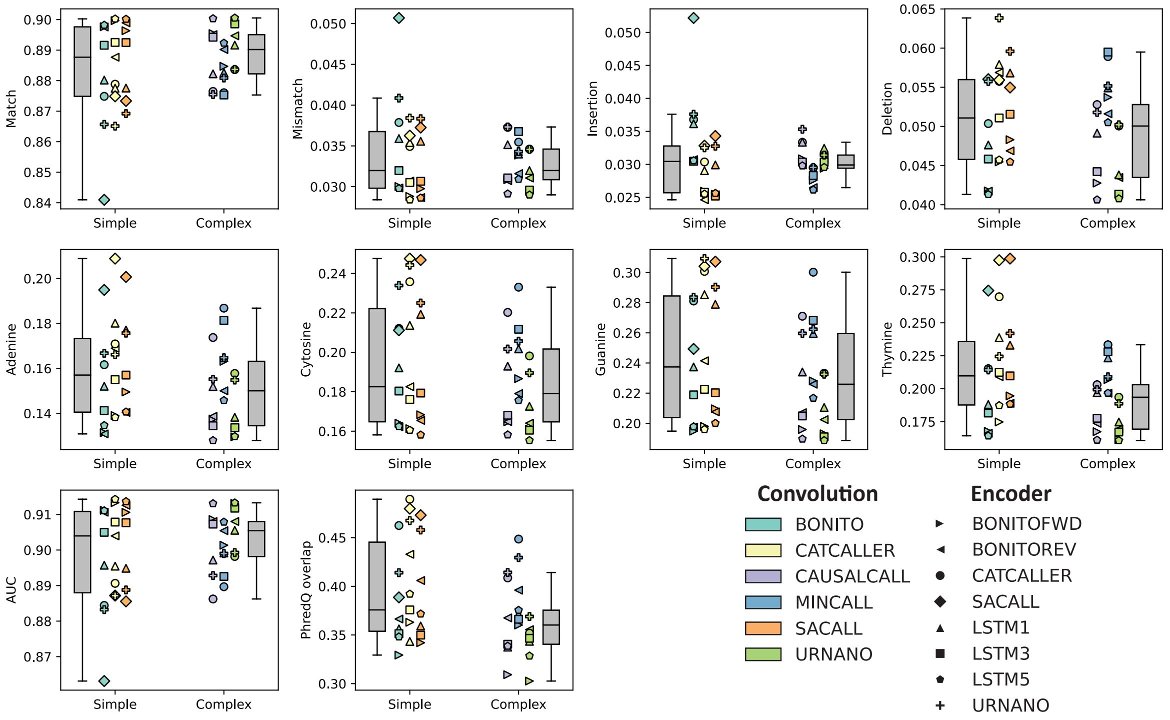

**Supplementary Figure 7: Comparison of simple and complex convolutions.** Performance comparison of CRF models that use a simple (*Bonito*, *CATCAller*, *SACall*) vs a complex (*Causalcall*, *Mincall*, *URNano*) convolution.

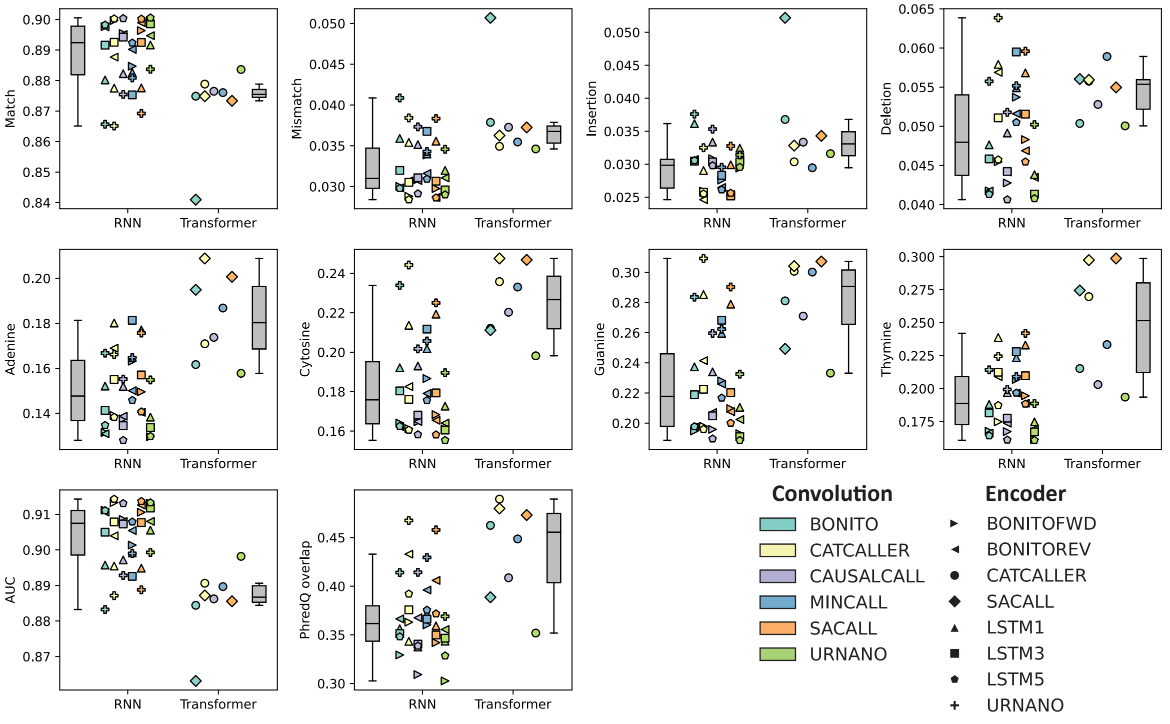

**Supplementary Figure 8: Comparison of RNN and Transformer encoders.** Performance comparison of CRF models that use a RNN encoder vs a Transformer encoder

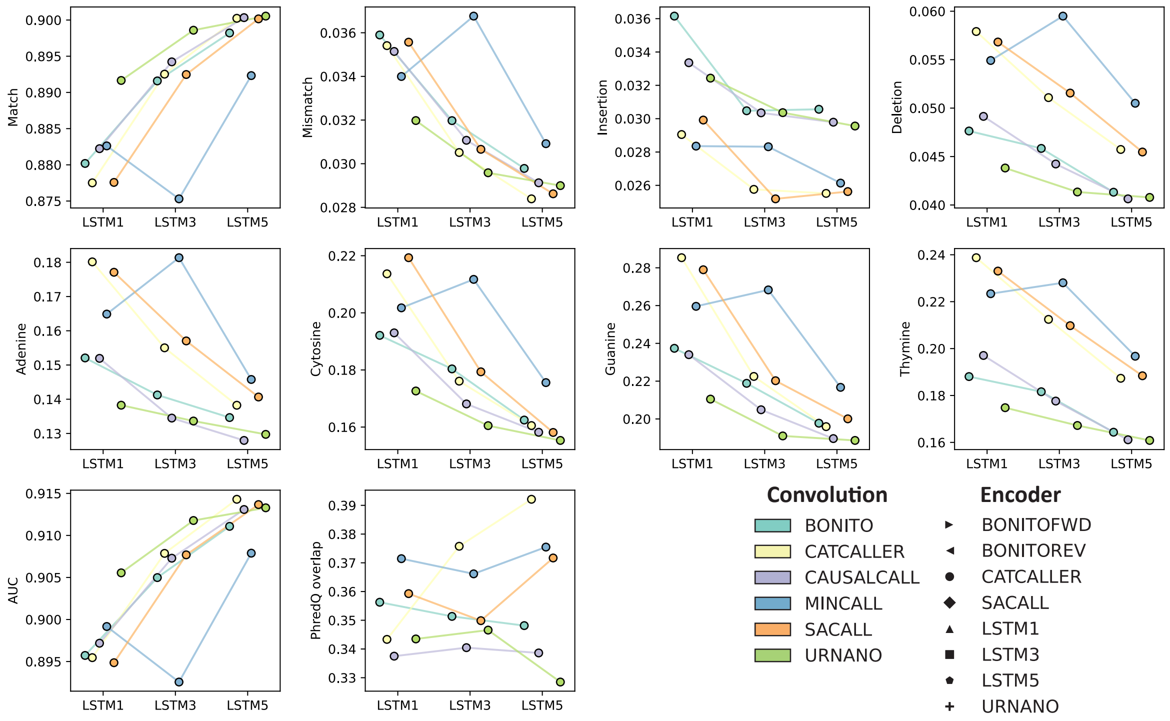

**Supplementary Figure 9: Comparison of LSTM detph.** Performance comparison of models with LSTM encoders with different layer depths (1, 3 or 5).

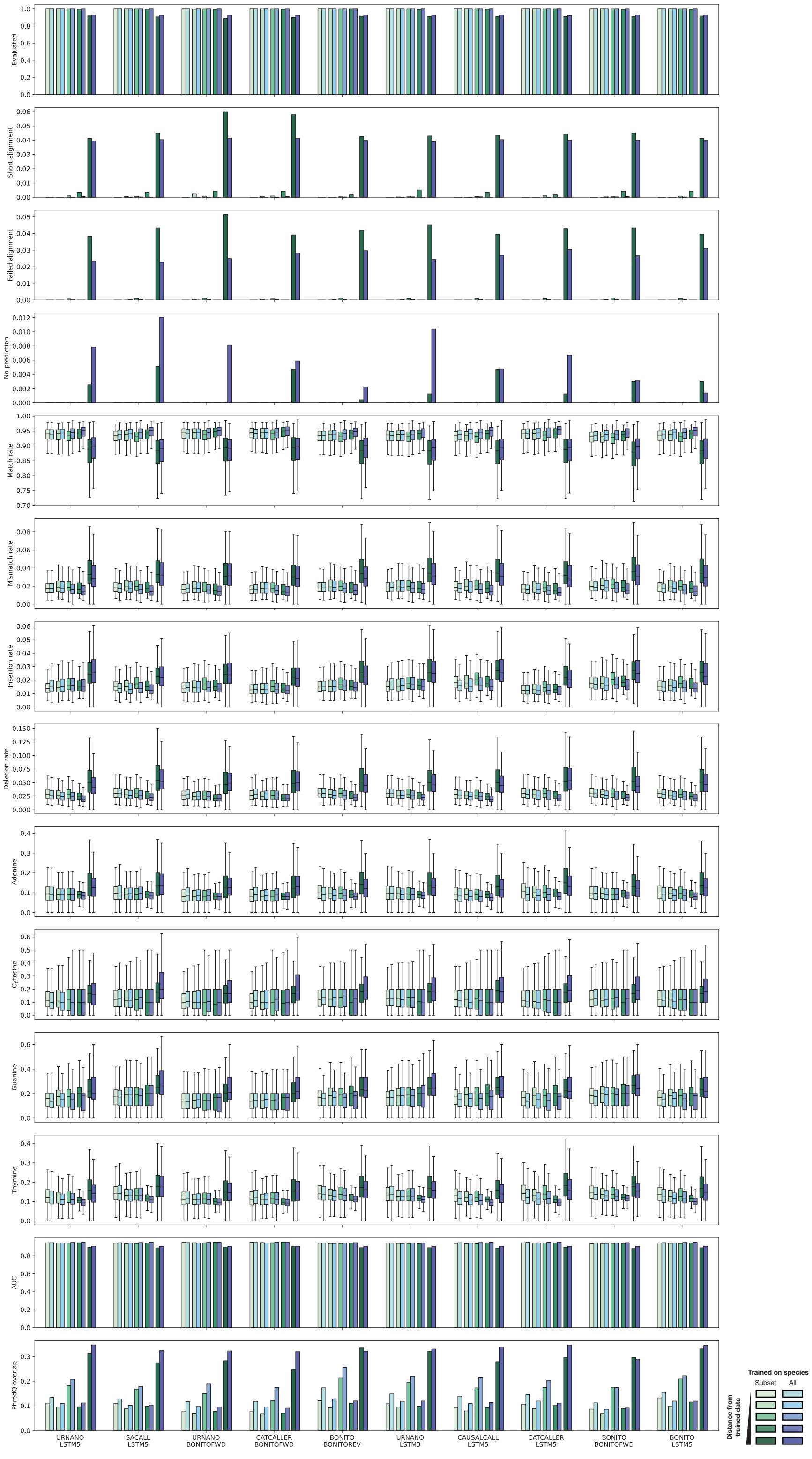

**Supplementary Figure 10: Task comparison of cross-species and global benchmarked models.** Comparison of the top 10 model combinations on trained on the cross-species or global datasets and tested on all species binned in the same manner as the cross-species dataset. Darker color indicates species are more different from the train set in the cross-species task.

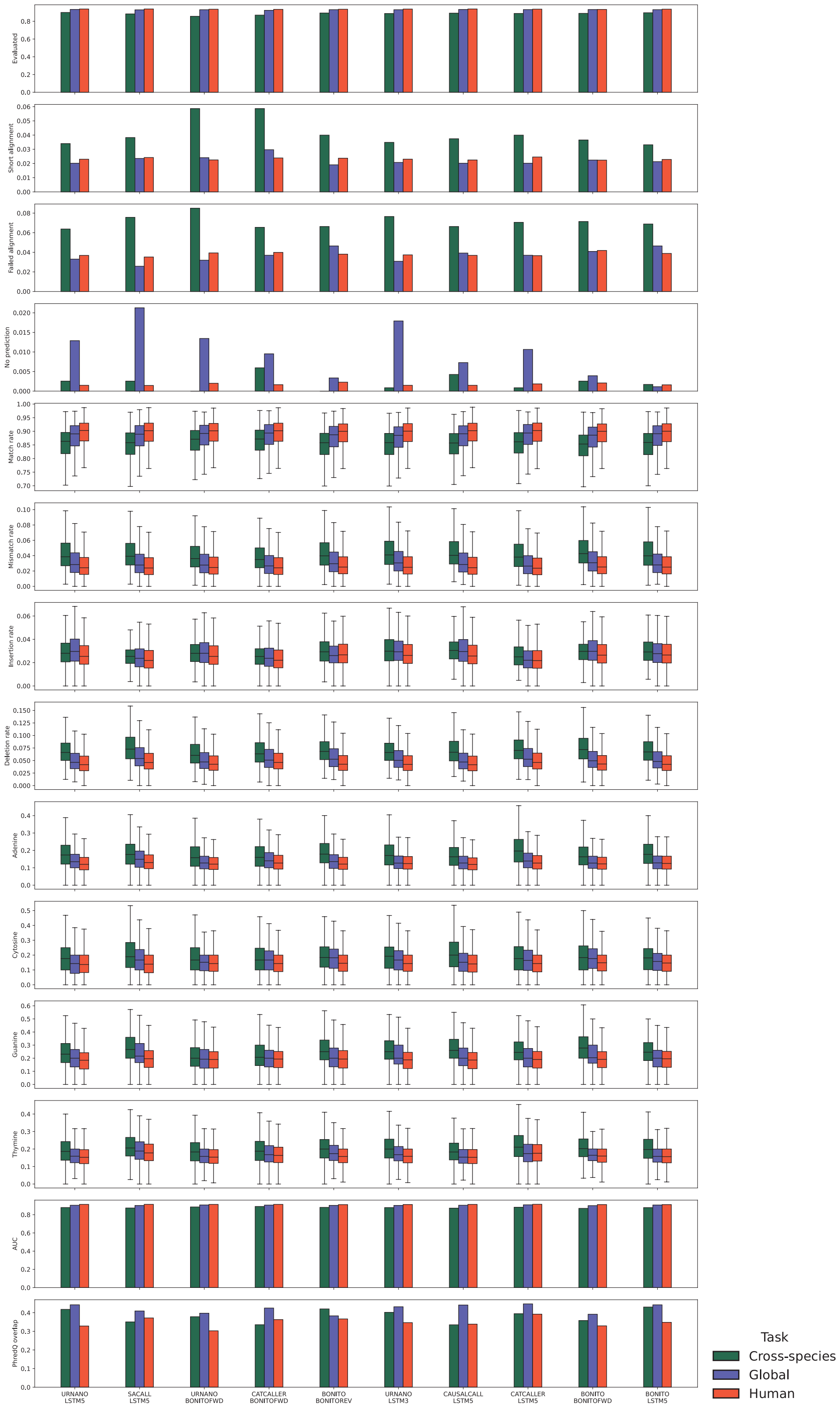

**Supplementary Figure 11: Task comparison of human, cross-species and global benchmarked models on human data.** Comparison of the top 10 model combinations on trained on the human, cross-species or global datasets and tested only on human data.

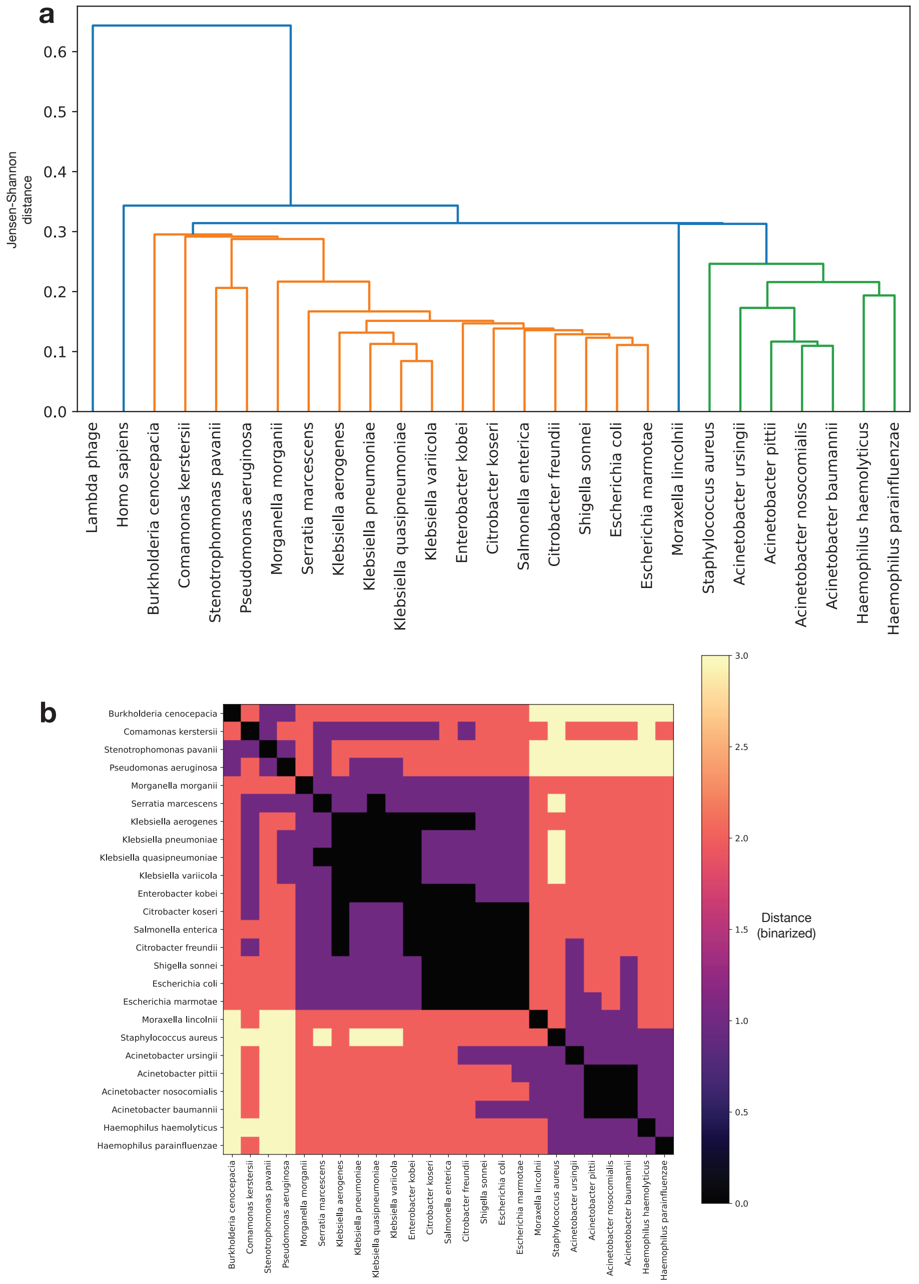

**Supplementary Figure 12: Clustering of benchmark dataset species.** (**a** Single-linkage hierarchical clustering of all the species based on the Jensen-Shannon divergence between the 9-mer relative counts of their genomes. **b** Heatmap with the binned distance between all bacterial species.

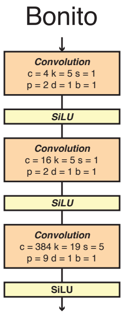

**Supplementary Figure 13: Convolutional architecture of Bonito.**

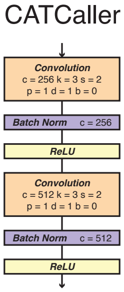

**Supplementary Figure 14: Convolutional architecture of CATCaller.**

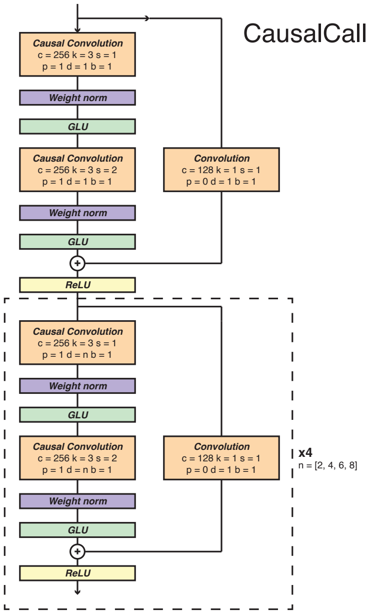

**Supplementary Figure 15: Convolutional architecture of CausalCall.**

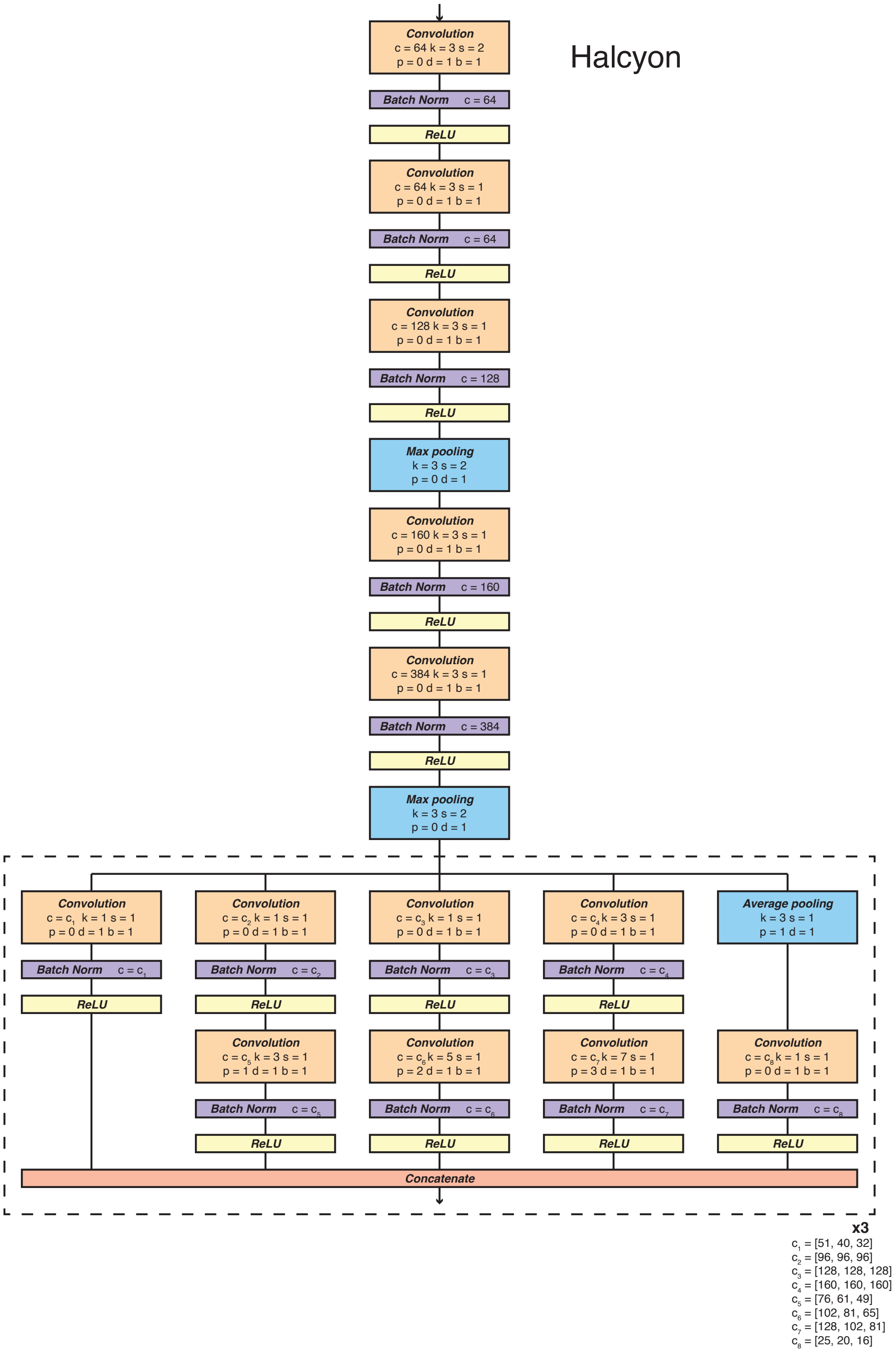

**Supplementary Figure 16: Convolutional architecture of Halcyon.**

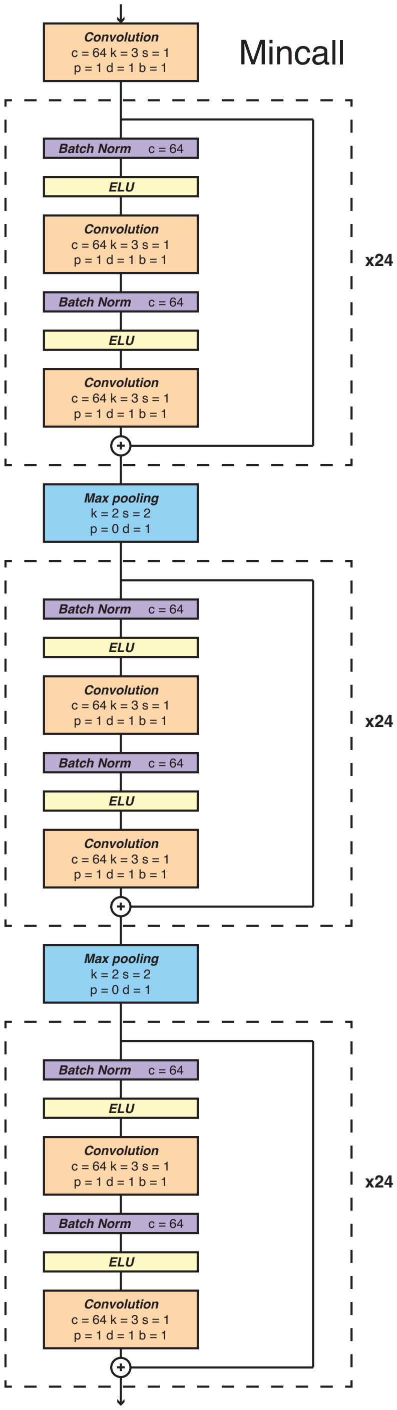

**Supplementary Figure 17: Convolutional architecture of Mincall.**

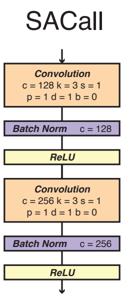

**Supplementary Figure 18: Convolutional architecture of SACall.**

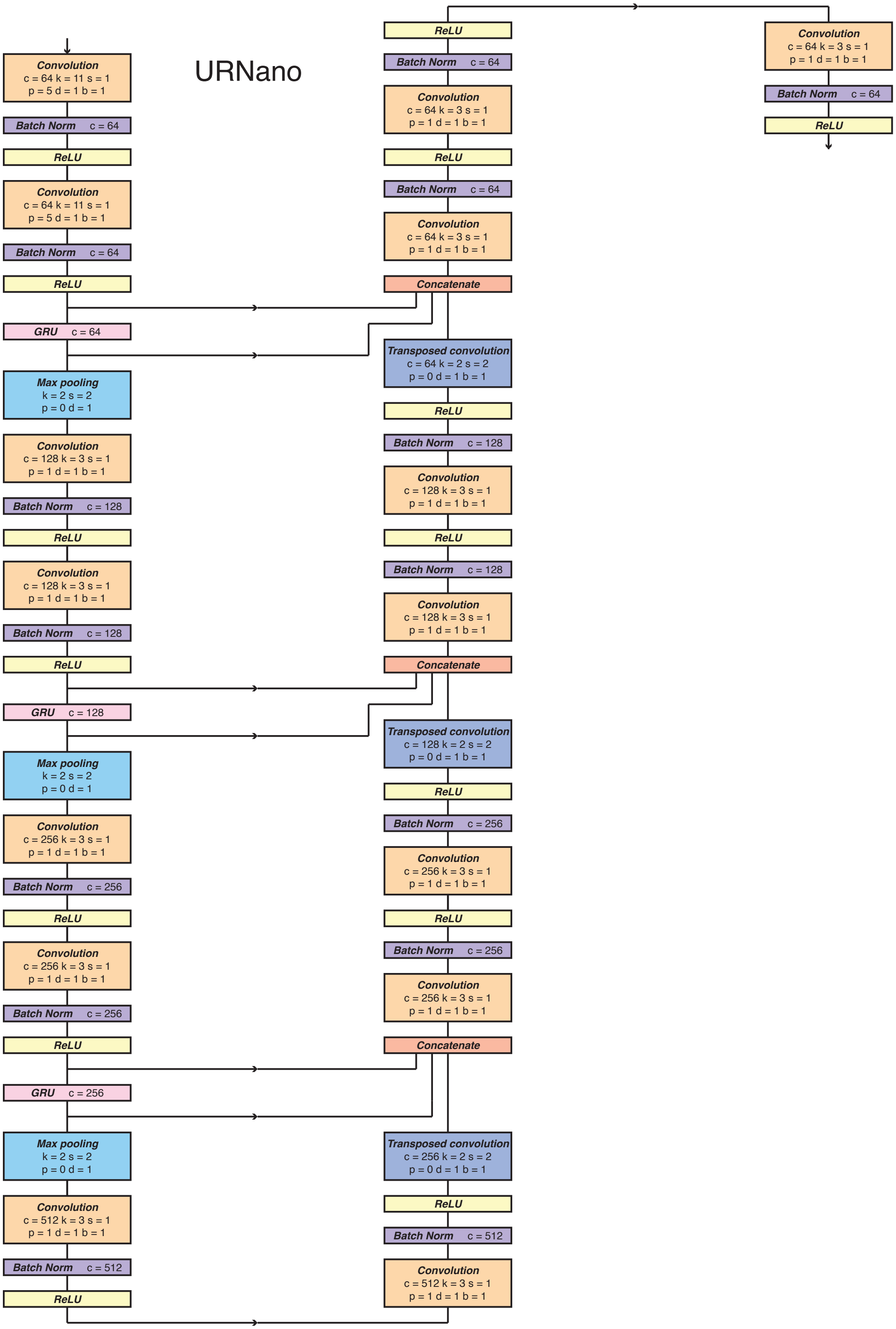

**Supplementary Figure 19: Convolutional architecture of URNano.**

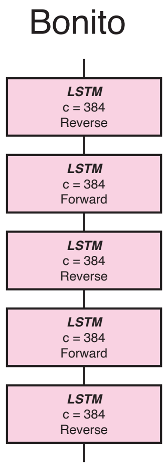

**Supplementary Figure 20: Encoder architecture of Bonito.**

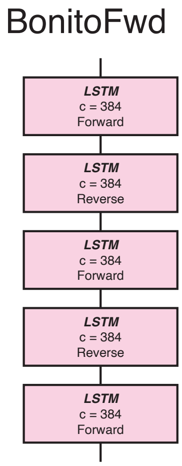

**Supplementary Figure 21: Encoder architecture of BonitoFwd.**

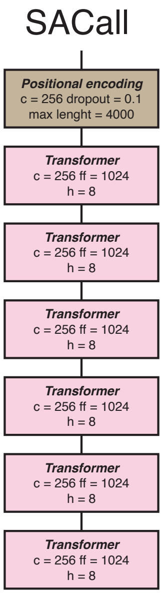

**Supplementary Figure 22: Encoder architecture of SACall.**

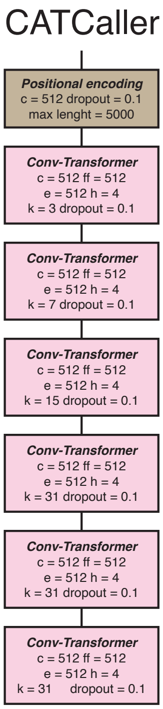

**Supplementary Figure 23: Encoder architecture of CATCaller.**

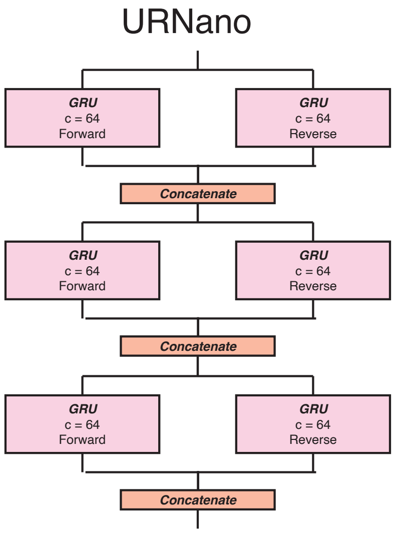

**Supplementary Figure 24: Encoder architecture of URNano.**

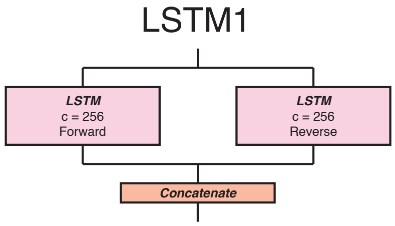

**Supplementary Figure 25: Encoder architecture of LSTM1.**

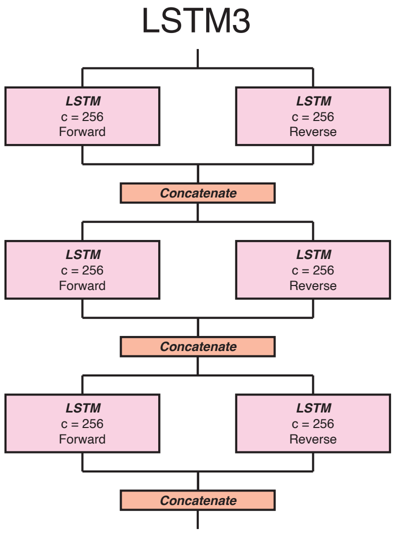

**Supplementary Figure 26: Encoder architecture of LSTM3.**

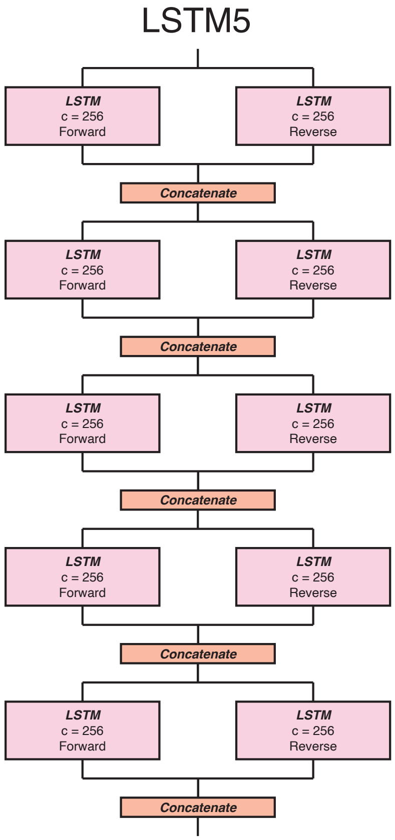

**Supplementary Figure 27: Encoder architecture of LSTM5.**

| Species | Dataset | Total reads | Benchmark reads | Global Train | Global Test | Human Train | Human Test | Cross-species Train | Cross-species Test |
|---|---|---|---|---|---|---|---|---|---|
| *Acinetobacter baumannii* | AYP-A2 | 6558 | 5768 | 3002 | 1785 | 0 | 0 | 0 | 1176 |
| *Acinetobacter nosocomialis* | MINF-5C | 6722 | 5992 | 3153 | 1785 | 0 | 0 | 0 | 1176 |
| *Acinetobacter pittii* | 16-377-0801 | 4467 | 4355 | 2193 | 1785 | 0 | 0 | 0 | 1176 |
| *Acinetobacter ursingii* | MINF-9C | 6976 | 6466 | 3069 | 1785 | 0 | 0 | 0 | 1176 |
| *Burkholderia cenocepacia* | MINF-4A | 7096 | 6401 | 3217 | 1785 | 0 | 0 | 0 | 1176 |
| *Citrobacter freundii* | MSB1-1H | 7093 | 6372 | 3244 | 1785 | 0 | 0 | 0 | 1176 |
| *Citrobacter koseri* | MINF-9D | 6900 | 6020 | 3003 | 1785 | 0 | 0 | 5000 | 500 |
| *Comamonas kerstersii* | MSB1-7G | 7242 | 6439 | 3446 | 1785 | 0 | 0 | 5000 | 500 |
| *Enterobacter kobei* | MSB1-1B | 7199 | 6291 | 3163 | 1785 | 0 | 0 | 5000 | 500 |
| *Escherichia coli* | MSB2-1A | 6985 | 6140 | 3157 | 1785 | 0 | 0 | 0 | 1176 |
| *Escherichia marmotae* | MSB1-5C | 7064 | 6431 | 3167 | 1785 | 0 | 0 | 0 | 1176 |
| *Haemophilus haemolyticus* | M1C132-1 | 8669 | 5984 | 2962 | 1785 | 0 | 0 | 0 | 1176 |
| *Haemophilus parainfluenzae* | M1C146-1 | 633 | 539 | 269 | 268 | 0 | 0 | 0 | 539 |
| *Homo sapiens* | FAB42828 | 33633 | 21666 | 862 | 398 | 9935 | 5836 | 0 | 287 |
| | FAF04090 | 94833 | 61778 | 2360 | 1206 | 28582 | 16803 | 0 | 792 |
| | FAF09968 | 21947 | 8942 | 349 | 181 | 4295 | 2361 | 0 | 97 |
| *Klebsiella aerogenes* | MINF-10B | 7200 | 6360 | 3150 | 1785 | 0 | 0 | 5000 | 500 |
| *Klebsiella pneumoniae* | INF007 | 1287 | 0 | 0 | 0 | 0 | 0 | 0 | 0 |
| | INF014 | 2449 | 0 | 0 | 0 | 0 | 0 | 0 | 0 |
| | INF032 | 15154 | 14353 | 0 | 0 | 0 | 0 | 834 | 91 |
| | INF042 | 11278 | 10141 | 0 | 0 | 0 | 0 | 583 | 53 |
| | INF065 | 1120 | 0 | 0 | 0 | 0 | 0 | 0 | 0 |
| | INF078 | 347 | 277 | 14 | 4 | 0 | 0 | 23 | 2 |
| | INF102 | 928 | 844 | 50 | 22 | 0 | 0 | 55 | 2 |
| | INF116 | 6775 | 0 | 0 | 0 | 0 | 0 | 0 | 0 |
| | INF125 | 1966 | 0 | 0 | 0 | 0 | 0 | 0 | 0 |
| | INF177 | 2853 | 0 | 0 | 0 | 0 | 0 | 0 | 0 |
| | INF192 | 2449 | 0 | 0 | 0 | 0 | 0 | 0 | 0 |
| | INF215 | 7142 | 6114 | 338 | 194 | 0 | 0 | 379 | 37 |
| | INF235 | 2282 | 0 | 0 | 0 | 0 | 0 | 0 | 0 |
| | INF310 | 3803 | 2892 | 166 | 105 | 0 | 0 | 175 | 19 |
| | INF319 | 2913 | 2127 | 118 | 45 | 0 | 0 | 131 | 9 |
| | INF321 | 1729 | 1374 | 91 | 41 | 0 | 0 | 85 | 7 |
| | INF322 | 7212 | 6446 | 351 | 189 | 0 | 0 | 350 | 44 |
| | INF341 | 1237 | 984 | 59 | 24 | 0 | 0 | 54 | 9 |
| | INF357 | 1032 | 843 | 46 | 18 | 0 | 0 | 60 | 1 |
| | INF358 | 2876 | 0 | 0 | 0 | 0 | 0 | 0 | 0 |
| | INF361 | 2628 | 2416 | 140 | 63 | 0 | 0 | 139 | 10 |
| | KSB1-1I | 7031 | 0 | 0 | 0 | 0 | 0 | 0 | 0 |
| | KSB1-6F | 245 | 198 | 13 | 10 | 0 | 0 | 5 | 0 |
| | KSB1-6G | 7040 | 5351 | 333 | 141 | 0 | 0 | 278 | 30 |
| | KSB1-7E | 5832 | 5540 | 297 | 177 | 0 | 0 | 288 | 32 |
| | KSB1-7F | 1636 | 0 | 0 | 0 | 0 | 0 | 0 | 0 |
| | KSB1-9A | 6787 | 5991 | 354 | 158 | 0 | 0 | 334 | 29 |
| | KSB1-9D | 4043 | 3624 | 187 | 106 | 0 | 0 | 207 | 19 |
| | KSB2-1B | 16847 | 0 | 0 | 0 | 0 | 0 | 0 | 0 |
| | NUH11 | 7336 | 6208 | 341 | 174 | 0 | 0 | 353 | 46 |
| | NUH27 | 7321 | 6169 | 389 | 149 | 0 | 0 | 367 | 36 |
| | KNUH29 | 15178 | 0 | 0 | 0 | 0 | 0 | 0 | 0 |
| | QMP-B2-170 | 459 | 395 | 28 | 10 | 0 | 0 | 17 | 1 |
| | SGH07 | 5645 | 4907 | 256 | 155 | 0 | 0 | 283 | 23 |
| *Klebsiella quasipneumoniae* | INF291 | 4047 | 3513 | 1812 | 1689 | 0 | 0 | 3013 | 500 |
| *Klebsiella variicola* | INF022 | 6501 | 6211 | 1810 | 877 | 0 | 0 | 2555 | 262 |
| | KSB1-8J | 6806 | 6039 | 1761 | 908 | 0 | 0 | 2445 | 238 |
| *Lambda phage* | VER5940 | 113514 | 111276 | 3571 | 1785 | 0 | 0 | 0 | 1176 |
| *Moraxella lincolnii* | 51409 | 1957 | 1715 | 731 | 865 | 0 | 0 | 0 | 1176 |
| *Morganella morganii* | MSB1-1E | 6307 | 5915 | 2866 | 1785 | 0 | 0 | 5000 | 500 |
| *Pseudomonas aeruginosa* | MINF-7A | 7082 | 6307 | 3106 | 1785 | 0 | 0 | 0 | 1176 |
| *Salmonella enterica* | 2010-06152 | 6638 | 6138 | 3142 | 1785 | 0 | 0 | 5000 | 500 |
| *Serratia marcescens* | 17-147-1671 | 16742 | 16262 | 3571 | 1785 | 0 | 0 | 5000 | 500 |
| *Shigella sonnei* | 2012-02037 | 9145 | 8711 | 3571 | 1785 | 0 | 0 | 0 | 1176 |
| *Staphylococcus aureus* | CAS38-02 | 11047 | 10858 | 3571 | 1785 | 0 | 0 | 0 | 1176 |
| *Stenotrophomonas maltophilia* | 17-G-0092-Kos | 16010 | 15075 | 3571 | 1785 | 0 | 0 | 0 | 0 |
| *Stenotrophomonas pavanii* | MSB1-4D | 3706 | 3067 | 1535 | 1426 | 0 | 0 | 0 | 1176 |
| - | **Total** | **615579** | **460225** | **81955** | **47088** | **42812** | **25000** | **48013** | **24355** |

**Supplementary Table 1: Summary of the collected datasets for benchmarking.** Collection of datasets used in this benchmark. Benchmark reads indicate the number of reads that could be aligned to their reference sequence. Number of reads used in each task for training and testing.

| Model | Basecalled reads (%) | | | | Alignment events (%) | | | | Hompolymer errors (%) | | | | PhredQ scoring | |
|---|---|---|---|---|---|---|---|---|---|---|---|---|---|---|
| | Pass | Failed mapping | Short alignment | No prediction | Match | Mismatch | Insertion | Deletion | Adenine | Cytosine | Guanine | Thymine | Overlap | AUC |
| *Bonito* | **93.4** | 4.1 | 2.3 | 0.2 | **90.0** | **2.5** | 2.6 | **4.3** | **12.5** | **14.9** | **19.7** | **15.8** | 32.4 | **0.910** |
| *CATCaller* | 92.4 | 5.2 | **2.2** | 0.2 | 86.5 | 3.6 | 3.6 | 5.9 | 14.2 | 23.5 | 27.4 | 18.2 | 8.2 | 0.886 |
| *Causalcall* | 66.3 | 25.7 | 7.8 | 0.2 | 77.6 | 6.0 | **1.7** | 14.4 | 43.4 | 60.0 | 50.7 | 42.0 | **0.7** | 0.802 |
| *Halcyon* | 79.5 | 4.7 | 15.4 | 0.4 | 81.3 | 3.3 | 3.2 | 8.8 | 20.8 | 20.0 | 22.0 | 25.8 | 11.7 | 0.844 |
| *Mincall* | 87.5 | 9.4 | 3.1 | **0.0004** | 83.7 | 4.9 | 3.7 | 7.2 | 17.2 | 21.8 | 28.6 | 20.0 | 6.6 | 0.863 |
| *SACall* | 92.2 | **3.7** | 2.4 | 1.7 | 86.5 | 3.5 | 3.0 | 6.5 | 20.0 | 22.6 | 29.6 | 19.7 | 8.7 | 0.886 |
| *URNano* | 90.4 | 4.8 | 3.3 | 1.6 | 85.4 | 3.6 | 2.0 | 8.6 | 28.8 | 28.6 | 35.6 | 32.6 | 7.2 | 0.879 |

**Supplementary Table 2: Original models benchmark summary.** Summary of the benchmark on training and testing the latest published basecallers on the human task. Results in bold denote best performance.

| Model | Basecalled reads (%) | | | | Alignment events (%) | | | | Hompolymer errors (%) | | | | PhredQ scoring | |
|---|---|---|---|---|---|---|---|---|---|---|---|---|---|---|
| | Pass | Failed mapping | Short alignment | No prediction | Match | Mismatch | Insertion | Deletion | Adenine | Cytosine | Guanine | Thymine | Overlap | AUC |
| *Bonito* | 93.4 | **4.1** | 2.3 | 0.2 | 90.0 | **2.5** | 2.6 | 4.3 | 12.5 | 14.9 | 19.7 | 15.8 | 32.4 | 0.910 |
| *Guppy original* | **95.2** | 4.4 | **0.4** | **0** | **91.3** | **2.5** | 2.0 | **3.9** | **9.7** | **14.7** | **18.9** | **12.5** | 27.9 | **0.937** |
| *Causalcall* | 66.3 | 25.7 | 7.8 | 0.2 | 77.6 | 6.0 | **1.7** | 14.4 | 43.4 | 60.0 | 50.7 | 42.0 | **0.7** | 0.802 |
| *Causalcall original* | 63.2 | 32.2 | 4.6 | **0** | 80.5 | 7.1 | 3.3 | 8.0 | 25.0 | 26.1 | 28.8 | 25.2 | 48.1 | 0.837 |
| *Halcyon* | 79.7 | 4.6 | 15.2 | 0.0004 | 81.1 | 3.3 | 3.2 | 8.7 | 20.8 | 20.0 | 22.0 | 25.9 | 11.7 | 0.844 |
| *Halcyon original* | 46.1 | 34.1 | 19.8 | **0** | 75.6 | 6.1 | 6.1 | 8.4 | 23.1 | 25.0 | 30.0 | 22.1 | N/A | N/A |

**Supplementary Table 3: Recreated models vs original models benchmark summary.** Comparison of existing basecallers and our recreations based on the human task benchmark. Results in bold denote best performance.

| Reference | Species | Dataset name | Pore version | Ligation kit | Available | Reference | Species | Dataset name | Pore version | Ligation kit | Available |
|---|---|---|---|---|---|---|---|---|---|---|---|
| [29] | *Klebsiella pneumoniae* | INF007 | R9.4 | - | Yes | [30] | *Homo sapiens* - NA12878 | FAB23716 | R9 | Rapid | Yes |
| | | INF014 | R9.4 | - | Yes | | | FAB39088 | R9.4 | Ligation | Yes |
| | | INF065 | R9.4 | - | Yes | | | FAB39075 | R9.4 | Ligation | Yes |
| | | INF078 | R9.4 | - | Yes | | | FAB39043 | R9.4 | Ligation | Yes |
| | | INF102 | R9.4.1 | - | Yes | | | FAB42706 | R9.4 | Ligation | Yes |
| | | INF116 | R9.4 | - | Yes | | | FAB41174 | R9.4 | Ligation | Yes |
| | | INF125 | R9.4 | - | Yes | | | FAB42260 | R9.4 | Ligation | Yes |
| | | INF177 | R9.4 | - | Yes | | | FAB42804 | R9.4 | Ligation | Yes |
| | | INF192 | R9.4 | - | Yes | | | FAB42316 | R9.4 | Ligation | Yes |
| | | INF215 | R9.4 | - | Yes | | | FAB42205 | R9.4 | Ligation | Yes |
| | | INF235 | R9.4 | - | Yes | | | FAB42561 | R9.4 | Ligation | Yes |
| | | INF310 | R9.4.1 | - | Yes | | | FAB42473 | R9.4 | Ligation | Yes |
| | | INF319 | R9.4.1 | - | Yes | | | FAB42395 | R9.4 | Ligation | Yes |
| | | INF321 | R9.4.1 | - | Yes | | | FAB42476 | R9.4 | Ligation | Yes |
| | | INF322 | R9.4 | - | Yes | | | FAB42451 | R9.4 | Ligation | Yes |
| | | INF341 | R9.4.1 | - | Yes | | | FAB42704 | R9.4 | Ligation | Yes |
| | | INF357 | R9.4.1 | - | Yes | | | FAB42828 | R9.4 | Ligation | Yes |
| | | INF358 | R9.4 | - | Yes | | | FAB42810 | R9.4 | Ligation | Yes |
| | | INF361 | R9.4.1 | - | Yes | | | FAB42798 | R9.4 | Ligation | Yes |
| | | KSB1-1I | R9.4 | - | Yes | | | FAB45280 | R9.4 | Ligation | Yes |
| | | KSB1-6F | R9.4 | - | Yes | | | FAB46664 | R9.4 | Ligation | Yes |
| | | KSB1-6G | R9.4 | - | Yes | | | FAB46683 | R9.4 | Ligation | Yes |
| | | KSB1-7E | R9.4 | - | Yes | | | FAB45332 | R9.4 | Ligation | Yes |
| | | KSB1-7F | R9.4 | - | Yes | | | FAB43577 | R9.4 | Ligation | Yes |
| | | KSB1-9A | R9.4 | - | Yes | | | FAB44989 | R9.4 | Ligation | Yes |
| | | KSB1-9D | R9.4 | - | Yes | | | FAF01169 | R9.4 | Ligation | Yes |
| | | NUH11 | R9.4 | - | Yes | | | FAF01441 | R9.4 | Ligation | Yes |
| | | NUH27 | R9.4 | - | Yes | | | FAB45277 | R9.4 | Ligation | Yes |
| | | QMP-B2-170 | R9.4 | - | Yes | | | FAB45321 | R9.4 | Ligation | Yes |
| | | SGH07 | R9.4 | - | Yes | | | FAF01127 | R9.4 | Ligation | Yes |
| | *Citrobacter freundii* | MSB1-1H | R9.4.1 | - | Yes | | | FAF01132 | R9.4 | Ligation | Yes |
| | *Citrobacter koseri* | MINF-9D | R9.4.1 | - | Yes | | | FAB49712 | R9.4 | Ligation | Yes |
| | *Enterobacter kobei* | MSB1-1B | R9.4.1 | - | Yes | | | FAF01253 | R9.4 | Ligation | Yes |
| | *Escherichia coli* | MSB2-1A | R9.4.1 | - | Yes | | | FAB45321 | R9.4 | Ligation | Yes |
| | *Escherichia marmotae* | MSB1-5C | R9.4.1 | - | Yes | | | FAB49914 | R9.4 | Ligation | Yes |
| | *Klebsiella aerogenes* | MINF-10B | R9.4.1 | - | Yes | | | FAB45271 | R9.4 | Ligation | Yes |
| | *Klebsiella quasipneumoniae* | INF291 | R9.4.1 | - | Yes | | | FAB49164 | R9.4 | Ligation | Yes |
| | *Klebsiella variicola* | INF022 | R9.4 | - | Yes | | | FAB49908 | R9.4 | Rapid | Yes |
| | | KSB1-8J | R9.4.1 | - | Yes | | | FAF04090 | R9.4 | Rapid | Yes |
| | *Salmonella enterica* | 2010-06152 | R9.4.1 | - | Yes | | | FAF15665 | R9.4 | Ultra | Yes |
| | *Acinetobacter baumannii* | AYP-A2 | R9.4 | - | Yes | | | FAF13748 | R9.4 | Ultra | Yes |
| | *Acinetobacter nosocomialis* | MINF-5C | R9.4.1 | - | Yes | | | FAF10039 | R9.4 | Ultra | Yes |
| | *Acinetobacter ursingii* | MINF-9C | R9.4.1 | - | Yes | | | FAF09968 | R9.4 | Ultra | Yes |
| | *Burkholderia cenocepacia* | MINF-4A | R9.4.1 | - | Yes | | | FAF09277 | R9.4 | Ultra | Yes |
| | *Comamonas kerstersii* | MSB1-7G | R9.4.1 | - | Yes | | | FAF14035 | R9.4 | Ultra | Yes |
| | *Haemophilus parainfluenzae* | M1C146-1 | R9.4.1 | - | Yes | | | FAF15694 | R9.4 | Ultra | Yes |
| | *Moraxella lincolnii* | 51409 | R9.4 | - | Yes | | | FAF09713 | R9.4 | Ultra | Yes |
| | *Morganella morganii* | MSB1-1E | R9.4.1 | - | Yes | | | FAF18554 | R9.4 | Rapid | Yes |
| | *Pseudomonas aeruginosa* | MINF-7A | R9.4.1 | - | Yes | | | FAF15630 | R9.4 | Ultra | Yes |
| | *Stenotrophomonas pavanii* | MSB1-4D | R9.4.1 | - | Yes | | | FAF09640 | R9.4 | Ultra | Yes |
| [5] | *Mycobacterium tuberculosis* | - | R9.4 | - | No | | | FAF09701 | R9.4 | Ultra | Yes |
| | *Phage lambda* | - | R9.4 | - | No | | | FAF15586 | R9.4 | Ultra | Yes |
| | *Escherichia coli* | - | R9.4 | - | No | | | FAF05869 | R9.4 | Ligation | Yes |
| [31] | *Escherichia coli* | - | R9 | - | Yes | This publication | *Lambda Phage* | VER5940 | R9.4.1 | Ligation | Yes |
| [17] | *Homo sapiens* - NA18943 | NA18943 | R9.4 | - | No | | | | | | |

**Supplementary Table 4: Datasets used for basecalling benchmarking.** List of publicly released Nanopore sequencing datasets that have been used for benchmarking.

| Species | Set | Difficulty |
|---|---|---|
| *Klebsiella variicola* | Train and test | 0 |
| *Serratia marcescens* | Train and test | 0 |
| *Klebsiella quasipneumoniae* | Train and test | 0 |
| *Comamonas kerstersii* | Train and test | 0 |
| *Citrobacter koseri* | Train and test | 0 |
| *Klebsiella pneumoniae* | Train and test | 0 |
| *Enterobacter kobei* | Train and test | 0 |
| *Salmonella enterica* | Train and test | 0 |
| *Klebsiella aerogenes* | Train and test | 0 |
| *Morganella morganii* | Train and test | 0 |
| *Citrobacter freundii* | Test | 1 |
| *Shigella sonnei* | Test | 1 |
| *Escherichia coli* | Test | 1 |
| *Escherichia marmotae* | Test | 1 |
| *Burkholderia cenocepacia* | Test | 2 |
| *Stenotrophomonas pavanii* | Test | 2 |
| *Pseudomonas aeruginosa* | Test | 2 |
| *Moraxella lincolnii* | Test | 2 |
| *Acinetobacter ursingii* | Test | 2 |
| *Acinetobacter pittii* | Test | 2 |
| *Acinetobacter nosocomialis* | Test | 2 |
| *Acinetobacter baumannii* | Test | 2 |
| *Haemophilus haemolyticus* | Test | 2 |
| *Haemophilus parainfluenzae* | Test | 2 |
| *Staphylococcus* aureus | Test | 3 |
| *Homo sapiens* | Test | 4 |
| *Lambda phage* | Test | 4 |

**Supplementary Table 5: Species split in the cross-species task.** Each species belongs to a bin depending on the k-mer genomic distance to the train-test species. Lower numbers indicate closer distance and higher numbers indicate further genomic distance.
